# Genome-wide phage susceptibility analysis in *Acinetobacter baumannii* reveals capsule modulation strategies that determine phage infectivity

**DOI:** 10.1101/2022.10.17.512463

**Authors:** Jinna Bai, Nicole Raustad, Jason Denoncourt, Tim van Opijnen, Edward Geisinger

## Abstract

Phage have gained renewed interest as an adjunctive treatment for life-threatening infections with the resistant nosocomial pathogen *Acinetobacter baumannii*. Our understanding of how *A. baumannii* defends against phage remains limited, although this information could lead to improved antimicrobial therapies. To address this problem, we identified genome-wide determinants of phage susceptibility in *A. baumannii* using Tn-seq. These studies focused on the lytic phage Loki, which targets *Acinetobacter* by unknown mechanisms. We identified 41 candidate loci that increase susceptibility to Loki when disrupted, and 10 that decrease susceptibility. Combined with spontaneous resistance mapping, our results support the model that Loki uses the K3 capsule as an essential receptor, and that capsule modulation provides *A. baumannii* with strategies to control vulnerability to phage. A key center of this control is transcriptional regulation of capsule synthesis and phage virulence by the global regulator BfmRS. Mutations hyperactivating BfmRS simultaneously increase capsule levels, Loki replication, and host killing, while BfmRS-inactivating mutations have the opposite effect, reducing capsule and blocking Loki infection. We identified novel BfmRS-activating mutations, including knockouts of a T2 RNase protein and the disulfide formation enzyme DsbA, that hypersensitize bacteria to phage challenge. We further found that mutation of a glycosyltransferase known to alter capsule structure and bacterial virulence can also cause complete phage resistance. Finally, additional factors including lipooligosaccharide and Lon protease act independently of capsule modulation to interfere with Loki infection. This work demonstrates that regulatory and structural modulation of capsule, known to alter *A. baumannii* virulence, is also a major determinant of susceptibility to phage.

**Author Summary:** Antibiotic-resistant infections with *Acinetobacter baumannii* are a major problem in critical care units and have increased in frequency during the COVID-19 pandemic. The virulence of these infections depends on a polysaccharide capsule surrounding the bacterium. Phage, or viruses that kill bacteria, represent a promising alternative therapy against highly antibiotic-resistant *A. baumannii* infections, and *A. baumannii*-specific phage often target the capsule. Here, we use high-throughput genetics to analyze how *A. baumannii* defends against phage and identify ways to potentiate their killing activity. We found that stressing the bacteria in ways that cause augmented production of capsule also causes hyper-susceptibility to phage. By contrast, turning off the stress response, or mutating the capsule structure, causes complete phage resistance. Altering another surface structure, lipooligosaccharide, or an intracellular protease also enhances phage attack. Modulating the amounts or makeup of capsular polysaccharide is known to influence virulence in *A. baumannii*. This work thus uncovers a connection between phage pressure and the evolution of virulence in *A. baumannii*, and it identifies control mechanisms that may be leveraged for improving future phage-based antimicrobial therapies.

## Introduction

The antibiotic-resistant, Gram-negative pathogen *Acinetobacter baumannii* is notorious for causing recalcitrant diseases in healthcare settings including sepsis and ventilator-associated pneumonia [1, 2]. Rates of *A. baumannii* infections resistant to last-line antibiotic treatments have increased during the COVID-19 pandemic [3, 4], and pan-resistant isolates have emerged [5, 6]. Therapy with lytic phages (viruses that infect and kill bacterial hosts) has gained renewed attention as an additional line of attack against infections with pathogens such as *A. baumannii* that are refractory to standard antibiotics. Phages have the benefit of having high host specificity, allowing selective targeting of pathogens without broad damage to the microbiome. Administration of anti-*A. baumannii* lytic phages in animal infection models has been efficacious in conferring protection from disease [4, 7–12]. Moreover, recent successes in the clinic have shown the promise of phage therapy as an adjunct to antibiotics for treating severe, progressive multidrug-resistant *A. baumannii* infections in critically ill patients [13–15]. Phage may also contribute to measures to block transmission of the pathogen in hospital environments [16].

Despite its potential, a drawback to phage therapy is the inevitable emergence of bacteria resistant to phage attack [17]. In the case of *A. baumannii*, this drawback may be mitigated by tradeoffs associated with phage resistance, such as lowered *in vivo* fitness/virulence, enhanced antibiotic sensitivity, or both [12, 17–19]. These costs likely emerge due to the *A. baumannii* capsule frequently serving as a requisite receptor for initiating phage infection [20–23].

Capsular polysaccharide is one of the most important virulence factors of *A. baumannii* [24–26]. Acapsular escape mutants isolated after phage treatment in animal models show dramatic attenuation similar to that documented for engineered capsule-null strains [12, 17, 18]. Although almost all *A. baumannii* patient isolates produce a capsule, the sugar compositions and structures of the capsular polysaccharide have diversified greatly across strains [27, 28]. These differences are in large part based on substitution of multiple genes within the highly variable K locus, which determines subunit sugar synthesis, polymerization, and export of the capsular polysaccharide [29, 30]. In some cases, structural changes can also manifest from variation in a single gene in the K locus [26, 31]. Different capsule structures may influence the interface with the host innate immune system, resulting in differences in how the capsule promotes pathogenicity [26, 32–34]. In addition to interstrain structural variation, *A. baumannii* varies the amount of capsule produced through regulatory networks. These capsule control networks include those that sense and respond to antimicrobial stress [35, 36] or that stochastically switch bacteria between two distinct phenotypic states [37–39], and each may contribute to how bacteria adjust to changing host or environmental conditions. The degree to which modulation of capsule levels affects susceptibility to phage attack is not known. Similarly, whether low-cost mutations modifying (rather than eliminating) capsule can provide avenues to phage resistance is also unclear. More broadly, functional characterization of *A. baumannii* determinants that facilitate or impede phage infection is largely lacking. Knowledge of these features could lead to strategies for improving the efficacy of anti-*A. baumannii* phage therapy and could enhance our understanding of the physiology and resilience of the organism.

Here, we use a functional genomic approach to investigate determinants of *A. baumannii*-phage interactions. We use Loki (vB_AbaS_Loki), a lytic phage of the *Siphoviridae* family that was isolated from activated sludge using *A. baumannii* ATCC 17978 [40]. Analysis of the Loki dsDNA genome has shown that it contains gene modules for virion morphogenesis, DNA replication, and lysis, as well as an uncharacterized MazG-family protein of unknown function [40]. The phage displays a narrow host range, infecting only 17978 among a panel of 40 diverse isolates, suggesting the use of a strain-specific receptor of unknown identity [40]. A component of the lipooligosaccharide (LOS) was suggested as the receptor, based on lack of Loki adsorption to an *lpxA* mutant [40]. In the present study, we have identified the primary Loki host receptor as the K3 capsule, and revealed pathways associated with capsule modulation that dramatically alter bacterial sensitivity to phage attack. This work has implications for phage therapy against *A. baumannii* and for understanding alteration of virulence in the pathogen.

## Results

### High-throughput identification of phage susceptibility determinants in *A. baumannii*

Loki infection of *A. baumannii* 17978 results in small, turbid plaques on solid medium and partial growth inhibition in liquid culture when using an initial multiplicity of infection (MOI) of 1 or 10 (Fig 1A) [40]. These features show that Loki has low virulence with 17978. We hypothesized that host mechanisms exist determining this outcome. To test this prediction and investigate genome-wide bacterial determinants of phage susceptibility, we used Tn-seq, taking advantage of our established strategies for identifying genetic determinants of antibiotic susceptibility in 17978 [41, 42]. Ten independent *Mariner* insertion pools each having about 20,000 individual insertions [43] (∼100,000 separate insertion sites in total) were grown for ∼8 doublings in parallel in broth containing Loki at different initial MOIs (1 to 1.5, or 10 to 15), or no phage (uninfected control). DNA was extracted from samples before (*t*_1_) and after (*t*_2_) the outgrowth of each pool. Massively parallel sequencing was used to measure the frequency of every transposon mutant in the population at each time point and calculate relative fitness (*W*) [41, 42] (see Materials and Methods). We then aggregated all the mutant fitness values (across all pools) associated with each gene to determine average, gene-level fitness scores (S1 Dataset). Genes were identified that had decreased or increased Tn-seq fitness during phage challenge compared to control, based on established criteria [fitness value derived from n ≥ 3 independent insertion mutants, fitness difference ≥ 10%, and false-discovery rate (FDR) ≤ 0.05 [41, 42]] (see Materials and Methods). These genes and their products are referred to as candidate phage susceptibility determinants, and those associated with decreased fitness upon phage challenge when mutated as candidate phage hypersusceptibility determinants.

**Fig 1.**
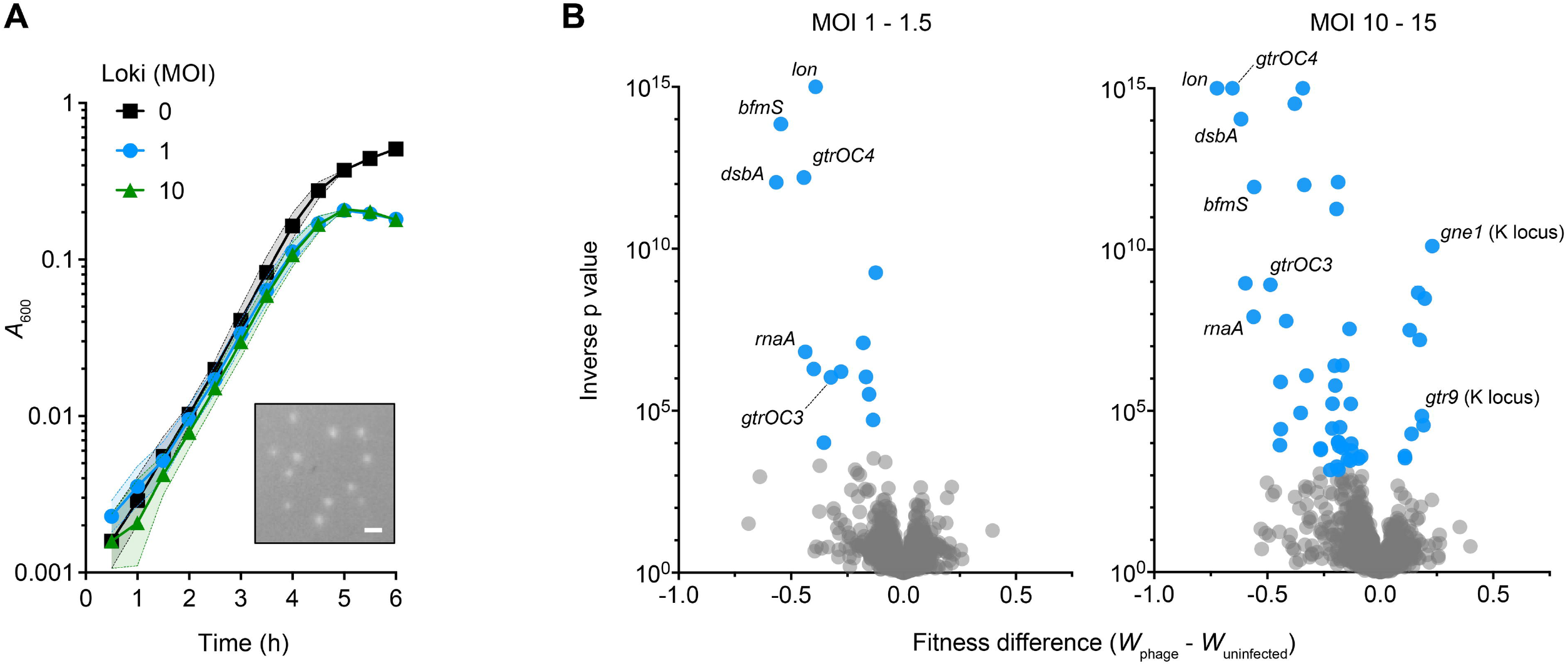
Identification of candidate determinants of susceptibility to Loki using Tn-seq. (A) Loki infection of *A. baumannii* 17978 causes partial inhibition of bulk culture growth. Bacteria were seeded in microplates with or without Loki at the indicated MOI and growth was monitored by OD readings (liquid infection assay). Data points show geometric mean ± s.d. (shaded bands) from n = 3 independent cultures. Where not visible, s.d. is within the confines of the symbol. Inset shows small, turbid plaques resulting from Loki infection of the same strain on solid medium. Plaques imaged with white light transillumination. Scale bar, 1 mm. (B) Tn-seq analysis of Loki susceptibility determinants in *A. baumannii* 17978. Volcano plots show change in gene-level fitness resulting from phage treatment at the indicated MOI compared to untreated control, plotted against inverse p value from parallel t tests. Blue data points are genes that pass significance criteria (Materials and Methods) and represent candidate Loki susceptibility determinants. Loci described further in the text are labelled. Grey data points are genes not passing significance criteria.

Challenge with Loki at the lower MOI (1 to 1.5) resulted in 14 candidate phage hypersusceptibility determinants (Fig 1B and S1 Dataset). The hits comprised numerous proteins involved in biosynthesis and control of the bacterial cell envelope. These included *bfmS*, the receptor of the BfmRS two-component system (TCS), which regulates capsule production, virulence, and resistance in response to signals that are yet to be defined [35, 36, 44]; LOS core oligosaccharide biosynthesis genes (*gtrOC2*, *gtrOC3*, *pda1*, *gtrOC4*, and *lpsB*) [29, 41]; and the *dsbA* dithiol oxidase [45] critical for forming protein disulfide bonds in the Gram-negative periplasm [46] (S1 Data Set). Additional hits were ACX60_RS02390 (referred to hereafter as *rnaA*), encoding a predicted periplasmic RNase T2 homolog; the *lon* protease; and the *dksA* transcription factor known to affect lytic phage infection in *E. coli* [47, 48].

Challenge with Loki at higher MOI (10 to 15) had two effects on the set of candidate susceptibility determinants identified. First, the number of phage hypersusceptibility determinants increased to 39 (Fig 1B and S1 Dataset). These hits included almost all (13/14) those identified using MOI 1-1.5, as well as additional enzymes belonging to the above functional categories (e.g., disulfide bond formation*, dsbC*, *dsbD*; LOS synthesis*, gtrOC8*; and the protease *clpA*) and to new functional categories (e.g., the DNA adenine methyltransferase, *aamA* [49]). Second, challenge with higher Loki MOI enabled detection of hits that had increased fitness (i.e., decreased susceptibility) compared to controls (Fig 1B and S1 Dataset). The strongest of these (showing greatest fitness increase and lowest q value) was *gne1*, which is a K locus gene required for generating precursors for capsule biosynthesis [29]. Loss of Gne1 completely prevents capsule production [50]. Mutations in an additional K locus gene, *gtr9*, which are also linked to capsule loss [19], also caused increased fitness during Loki challenge (S1 Dataset). The other hits were extrinsic to the K locus, but several had annotations related to polysaccharide production. The Tn-seq screen thus reveals a variety of candidate factors that increase or decrease *A. baumannii* vulnerability to phage attack.

### Targeted deletion and complementation analyses validate the phage hypersusceptibility phenotypes predicted by Tn-seq

We confirmed the Tn-seq results using individual mutant strains, focusing first on candidate phage susceptibility determinants from different functional categories showing lower Tn-seq fitness with Loki challenge (Fig 2A). Strains carrying in-frame deletions of *rnaA*, *dsbA*, *gtrOC3*/*4*, and *lon* were constructed (Materials and Methods). To enable *trans* complementation analysis, plasmids harboring an IPTG-inducible WT allele, or a control plasmid lacking the gene, were introduced into the respective mutant strains (S1 Table). In the case of *bfmS*, in lieu of complementation analysis we took advantage of multiple different mutants we previously isolated in the *bfmRS* locus. This includes two *bfmS* mutants—a deletion with *aacC1* replacement (referred to as Δ*bfmS*), and the frame-shift G467Dfs*19 [35] (referred to as *bfmS*\*)—as well as additional mutants affecting *bfmR* as will be explained in a subsequent section. Susceptibility to Loki was assessed by liquid challenge and plaque formation assays (Materials and Methods). During liquid phage challenge, the *rnaA*, *dsbA*, and *bfmS* mutants all showed complete block of growth immediately after exposure to Loki, unlike the WT, which showed only partial growth inhibition after about 5 hours (Fig 2B and C). Different kinetics were seen with Loki challenge of the other mutants. With the Δ*gtrOC3* and Δ*gtrOC4* mutants exposed to phage, bacteria grew for an initial ∼2.5 hours before collapse (Fig 2B). With Δ*lon*, by contrast, Loki inhibited bacterial growth until ∼4 hours, at which point rapid growth resumed (Fig 2B). Reintroduction of the respective WT genes reversed the Loki hypersusceptibility phenotypes of *rnaA, dsbA*, *gtrOC3*, *gtrOC4*, and *lon* mutants. In some cases, leaky expression without IPTG was sufficient for partial (*dsbA*, *gtrOC4*) or full (*gtrOC3, lon*) complementation (Fig 2B). Addition of IPTG at 0.1 mM allowed reversal of the Loki hypersusceptibility phenotype of all strains, although a partial growth delay defect remained with the *rnaA* reintroduction strain exposed to phage (Fig 2B). Raising the level of *rnaA* induction with 1 mM IPTG completely eliminated this defect and had the added effect of rendering the strain even less susceptible to Loki than the WT control (S1A Fig).

**Fig 2.**
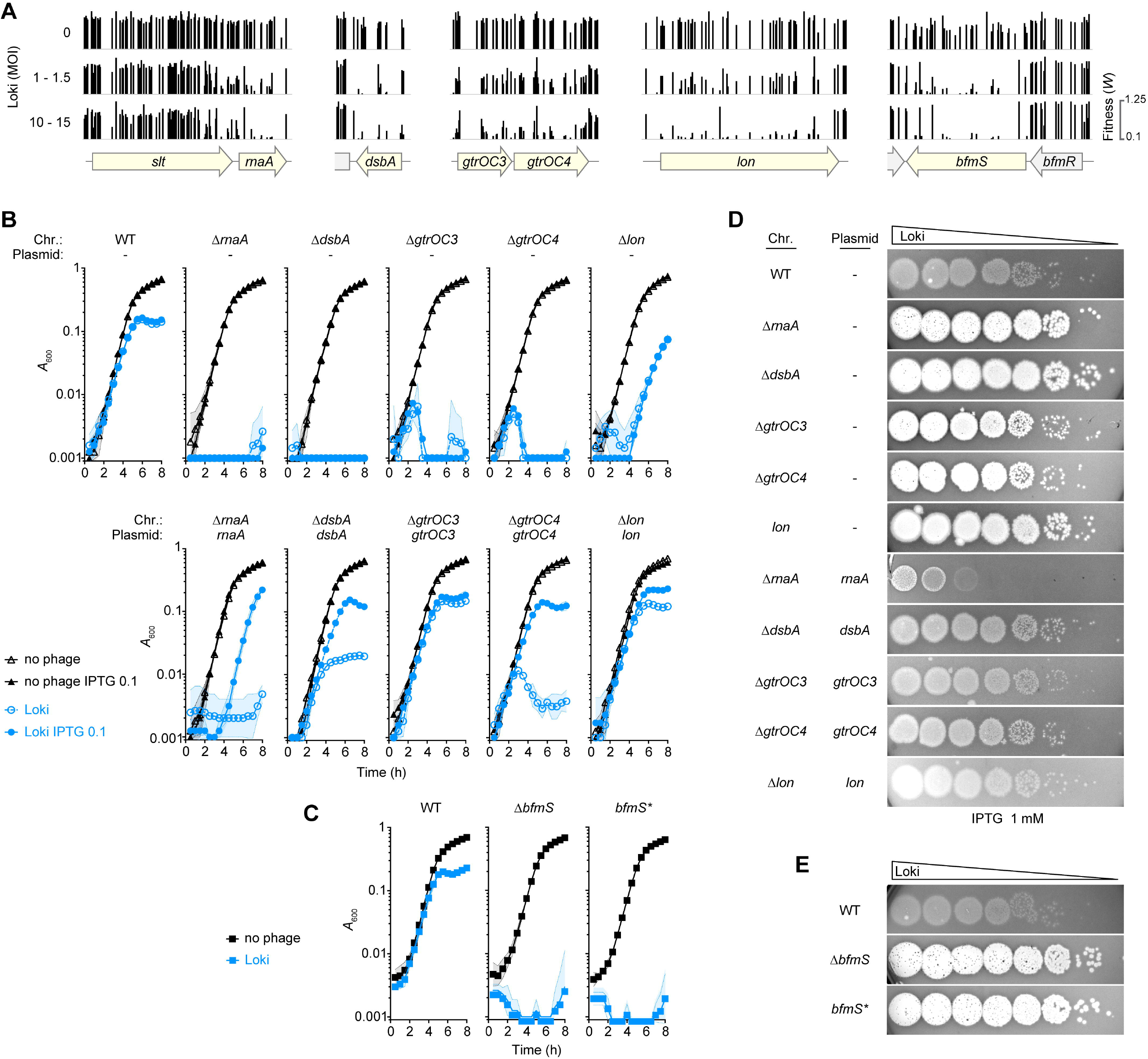
Validation of candidate phage hypersusceptibility loci. (A) Tracks show Tn-seq fitness values (vertical bars) of individual *Mariner* transposon mutants at candidate Loki hypersusceptibility loci (shaded yellow) across all tested mutant banks using the indicated MOI. (B-C) Liquid challenge assays with defined mutants. Growth of the indicated *A. baumannii* 17978 strains with Loki (MOI 1) or no phage was measured as in Fig 1A (n=3). “Chr” indicates the chromosomal mutation; “Plasmid” denotes the gene reintroduced under P(IPTG) control via vector pYDE152; - indicates control plasmid with no reintroduced gene; *bfmS*\* refers to *bfmS*-null point mutant EGA127; IPTG “0.1” and “1” refer to mM units. (D-E) Plaque formation assays. ∼10^8^ of the indicated bacteria were overlaid on bottom agar, followed by spotting 10-fold serial dilutions of Loki. Bottom agar in D contained 1mM IPTG. Plaques were imaged with white-light transillumination after 16 h incubation at 30°C.

Results from plaque formation assays mirrored the findings from the liquid challenge experiments. In contrast to the small, turbid plaques caused by Loki with control bacteria, the same phage caused large, clear plaques with each candidate hypersusceptibility mutant (Fig 2D and E). Plaques appeared to be larger with *rnaA, dsbA*, and *bfmS* mutants compared to *gtrOC3* and *gtrOC4*, echoing the susceptibility differences seen with Loki challenge in broth (Fig 2, compare D, E to B, C). Expressing the respective genes *in trans* with 1 mM IPTG reversed the enhanced plaque phenotypes in all cases (Fig 2D). In particular, *rnaA* overexpression greatly lowered the plaque formation efficiency of Δ*rnaA* beyond the level of the WT control, similar to its effect during liquid challenge (Fig 2D and S1A Fig, compare Δ*rnaA*/*rnaA* to the WT control).

We note that the gene directly upstream of *rnaA*, ACX60_RS02385 (*slt* [36]), was also identified as a candidate susceptibility determinant associated with modestly decreased fitness during Loki challenge (S1 Dataset). Exposure of an in-frame *Δslt* mutant to Loki recapitulated the modest hypersusceptibility defect (S2A, B Fig). Analysis of the *slt* Tn-seq mutant fitness values revealed a potentially biased distribution, with insertions causing major Loki hypersusceptibility defects having an apparent bias toward the 3’ end of the locus that is proximal to *rnaA* (Fig 2A). The Δ*slt* mutant also showed slightly reduced levels of *rnaA* expression (S2C Fig). Since phage susceptibility appears sensitive to *rnaA* expression level (Fig 2B, D, E), we cannot rule out a polar effect on *rnaA* in the altered phage susceptibility of the *slt* mutant, and this mutant was not pursued further. These results underscore that candidate Loki hypersusceptibility loci identified through Tn-seq show the predicted susceptibility phenotypes when analyzed independently using engineered mutant strains.

### Analysis of Loki resistance mutations reveals capsular polysaccharide as phage receptor

Given the identification of capsule production genes as sites of mutations that increase fitness during phage challenge (S1 Dataset), we tested the hypothesis that the capsule is essential to the infective cycle of Loki by serving as a host receptor. To this end, we examined the Tn-seq data associated with the entire K locus of 17978 (KL3) and additional genes involved in capsule production. Gene-level Tn-seq fitness scores were available for 9 out of 20 KL3 genes. The other 11 were represented by too few transposon mutants, as observed previously, likely due to their essential involvement in late steps of capsule biosynthesis that may cause dead-end intermediates when blocked [43]. For most of the 9 permissive KL3 genes, transposon disruption led to higher fitness during Loki exposure relative to untreated control (with 5 having non-FDR P < 0.05; Fig 3A). Disruption of *wzi*, an additional permissive gene outside the K locus that encodes the lectin required for efficient surface binding of capsular polysaccharide [51], also increased Tn-seq fitness during Loki challenge compared to control, albeit without passing statistical cut-offs (P = 0.15). Since a subset of the glycan building blocks encoded by the K locus are used for protein glycosylation by PglL [52], we also examined the *pglL* Tn-seq phenotypes. In contrast to the capsule synthesis genes, transposon insertions in *pglL* showed slightly decreased fitness during Loki challenge (Fig 3A). These findings indicate that lesions leading to capsule deficiency are associated with enhanced fitness in the face of Loki challenge, while the opposite is true for *pglL* mutations, which are known to eliminate protein O-glycans but preserve capsule [52].

**Fig 3.**
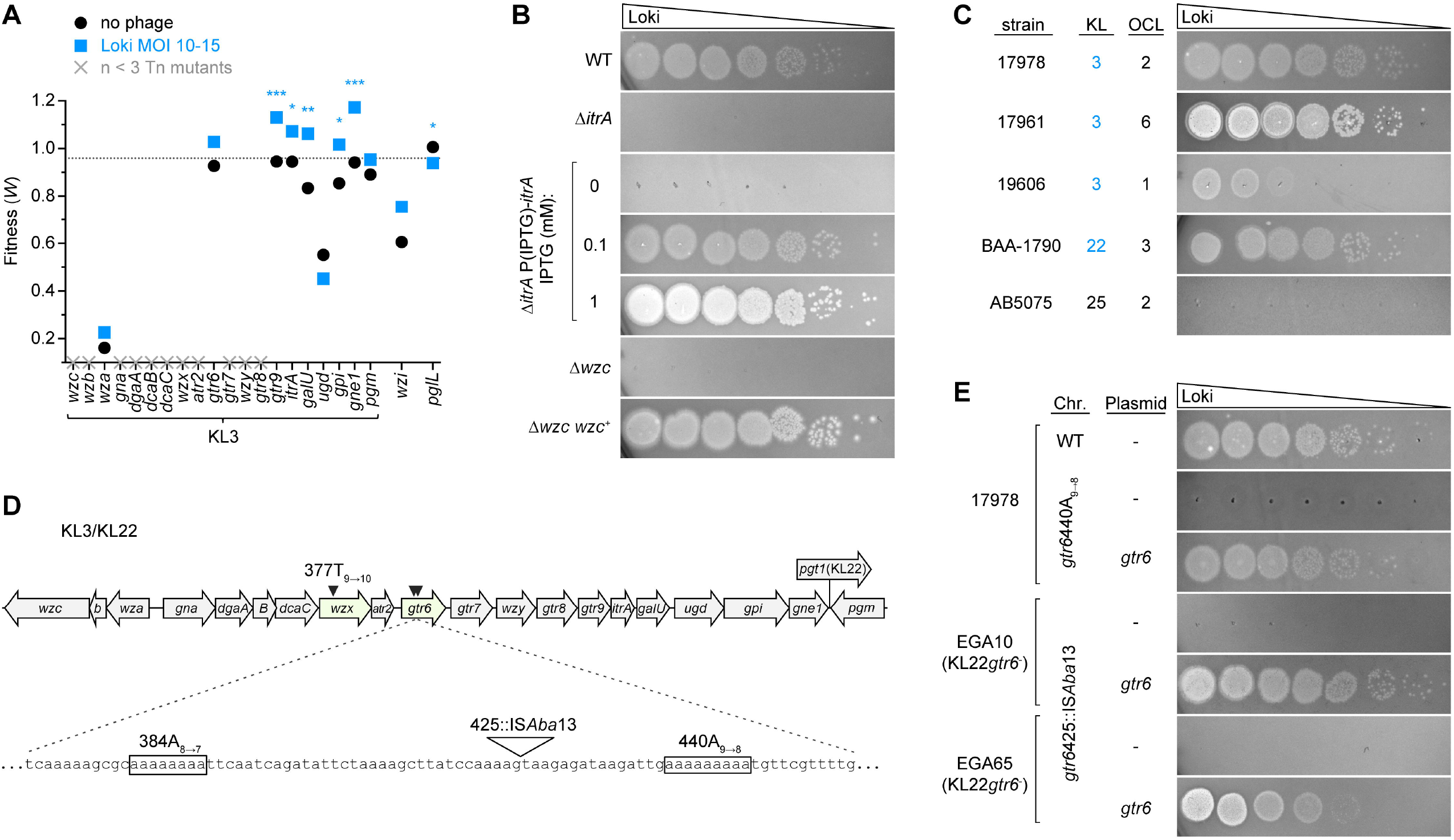
Loki resistance mutations map to capsule biosynthesis locus. (A) Tn-seq fitness of transposon mutants in the capsule synthesis (KL3) and associated loci in the Loki challenge screen. Plotted are the average gene-level Tn-seq fitness values (*W*) with phage infection (Loki MOI 10-15) vs uninfected control, analyzed by unpaired t-tests. X indicates essential gene without a fitness value due to paucity of transposon insertions. Dotted line indicates the genome-wide average *W* during Loki challenge. (B-C, E) Plaque formation assays showing dependence on the KL3 capsule for Loki infection. Loki was spotted on the following hosts: (B) acapsular *itrA* and *wzc* mutants of 17978; (C) a range of *A. baumannii* clinical isolates harboring the indicated K and OC locus (K loci producing the K3 capsule type are shaded blue); or (E) strains with spontaneous *gtr6* mutations (Table S1). Strains in E harbored the indicated pYDE152-based plasmid and bottom agar contained 1mM IPTG. (D) Diagram of the KL3/KL22 cluster showing spontaneous Loki resistance mutations (arrowheads). The hotspot within *gtr6* is expanded. KL3 and KL22 share all ORFs except *pgt1* (only present in KL22).

Building on these observations, we determined the requirement of capsule for Loki infection by examining (1) targeted capsule mutants, (2) additional clinical isolates of diverse capsule types, and (3) spontaneous phage-resistant mutants. First, we analyzed deletions in 17978 of two critical capsule synthesis genes: *itrA*, the initiating transferase required for use of phosphosugars in both capsule synthesis and protein O-glycosylation and *wzc*, essential to assembly and export of high-molecular weight capsular polysaccharide [29, 35, 52]. Δ*itrA* causes loss of both capsule and protein O-glycans, while Δ*wzc* blocks capsule specifically. We note that unlike *itrA*, *wzc* is essential in 17978 [35, 43]. Δ*wzc* could be analyzed, however, by the parallel use of EGA106-27, which is a viable 17978 Δ*wzc* isolate harboring extragenic compensatory mutations, and EGA260, its isogenic *wzc*-repaired derivative obtained by marker rescue [35]. As predicted by Tn-seq, deletion of *itrA* led to increased growth during Loki challenge in liquid (S3A Fig) and completely blocked plaque formation (Fig 3B). Reintroduction of *itrA* reversed this defect, with the resulting susceptibility to Loki dependent on the level of *itrA* induction (Fig 3B; S3A Fig). Loki was also completely unable to form plaques on EGA106-27 (Wzc^-^), but readily infected EGA260 (Wzc^+^) (Fig. 3B). The plaques on EGA260 were larger and clearer than that seen with the WT control, presumably a consequence of the compensatory mutations rendering it hypersensitive to phage. The Loki non-susceptibility of ItrA-deficient (capsule^-^/protein O-glycan^-^) and Wzc-deficient (capsule^-^/protein O-glycan^+^) strains, and the hypersusceptibility detected with *pglL* transposon mutants (capsule^+^/protein O-glycan^-^), points to capsular polysaccharide as essential for phage infection.

Second, we found that the K3 capsule type correlates with the ability of diverse clinical isolates to be targeted by Loki. To determine Loki host range we used strains 17978, 17961 and 19606, which all have KL3; BAA-1790 [53], which has KL22 (identical to KL3 but with one extra gene; Fig. 3D); and AB5075, which has the unrelated KL25 cluster. Note that both KL3 and KL22 biosynthetic gene clusters yield the same capsular polysaccharide, type K3 [27, 54], while the K25 capsule generated from KL25 has a completely different structure [55]. Loki formed plaques on all four K3 strains, but not the K25 strain (Fig. 3C). Ability to infect was independent of the type of LOS outer core biosynthesis locus (OCL) in the host strains (Fig. 3C). These results are consistent with the K3 capsule, and not LOS outer core, providing a specific host receptor for Loki infection.

Third, analysis of spontaneous Loki-resistant mutants supported the model that capsule provides an essential, specific receptor for Loki. Three independent 17978 mutants forming colonies within zones of confluent lysis were isolated and their genomes were sequenced (Materials and Methods). Each strain had a single, slipped-strand mispairing mutation within homopolymeric sites in the K locus (Fig 3D). Two were identical mutations (440A_9→8_) in *gtr6*, encoding a glycosyltransferase that adds a single sugar side-chain to the capsule repeat unit [26] (S2 Table), and the third was in the *wzx* polysaccharide flippase (377T_9→10_) [29]. Each mutation caused a frameshift and premature translation termination. While deficiency of Wzx is expected to result in complete capsule loss, Gtr6 deficiency is known to preserve capsule but cause an altered structure lacking the side chain sugar [26]. Intriguingly, *gtr6* deletion was recently shown to amplify 17978 virulence during bloodstream infection in mice, and a naturally occurring *gtr6* mutation (1008::IS*Aba13*) was found to drive hypervirulence in another K3 isolate, HUMC1 [26]. We confirmed that 17978 *gtr6*440A_9→8_ resists Loki plaque formation, a phenotype reversed by reintroduction of WT *gtr6* (Fig 3E) and recapitulated by an in-frame *gtr6* deletion (S3B Fig). Consistent with previous findings that loss of Gtr6 alters capsule structure but not amount [26], *gtr6*440A_9→8_ and Δ*gtr6* caused no change to overall capsule levels (S4A Fig). Search of public databases revealed that the 440A_9→8_ allele is present in a variety of KL3 and KL22 *A. baumannii* clinical isolates (S2 Table and ref. [56]), consistent with this site as a mutational hotspot in *gtr6*. We also examined *gtr6* within two strains harboring KL22 from our sequenced *A. baumannii* clinical isolate collection, EGA10 and EGA65. We found that both strains have a novel IS*Aba13* insertion near 440A (425::IS*Aba*13), resembling the situation with HUMC1 [26]. EGA10 and EGA65 were both Loki-resistant, but introducing WT *gtr6* from 17978 had the dramatic effect of conferring Loki susceptibility with efficient plaque formation (Fig 3E). These results further support the K3 capsule as a specific Loki receptor, and suggest that *gtr6* is a hub for mutations linking phage evasion to the evolution of *A. baumannii* virulence.

Finally, we examined the effect of *itrA*, *gtr6*, and *wzc* mutations on phage adsorption (Materials and Methods). Each mutation resulted in loss of Loki attachment, as indicated by high fractions of phage remaining in suspension after mixture with bacteria (Fig 4A). Together these results support the conclusion that Loki uses the K3 capsule as an essential receptor.

**Fig 4.**
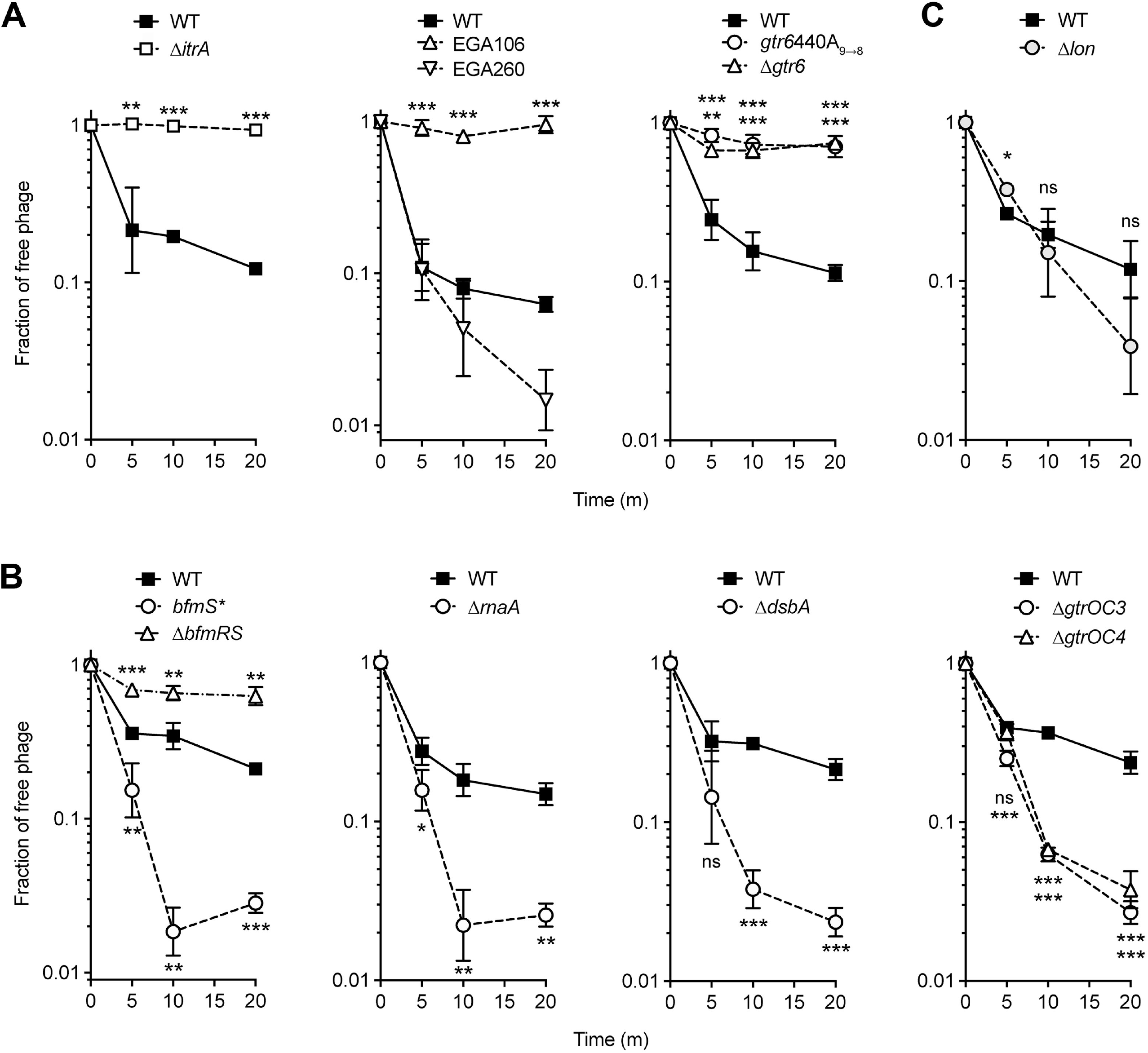
Results of phage adsorption assays measuring *A. baumannii* binding to Loki. Loki adsorption assays were performed with WT 17978 or the indicated mutant. Data points show geometric mean ± s.d. (n=3), analyzed by unpaired t test (mutant vs WT control). P values: *, ≤0.05; **, ≤0.01; ***, <0.001; ns, not significant.

### BfmRS activity controls phage susceptibility

Loki-hypersensitizing *bfmS* mutations lead to capsule hyperproduction by increasing transcription of capsule biosynthesis genes [35], likely through enhanced activity of the cognate transcription factor BfmR [44]. Controlling BfmR activity and capsule production could thereby also allow control of *A. baumannii* susceptibility to phage. We tested this model by analyzing mutations in *bfmR*, which cause phenotypes opposite those of *bfmS* mutants including decreased capsule production [35]. *bfmR* mutants are thus predicted to be less susceptible to capsule-targeting phage. In the Tn-seq screen, *bfmR* transposon mutants indeed had increased fitness during Loki challenge relative to controls, particularly at the higher MOIs, in stark contrast to the dramatic hypersusceptibility of *bfmS* mutants (Fig 5A). To confirm the altered phage susceptibility phenotype associated with *bfmR*, we used a knockout of the entire TCS (Δ*bfmRS*), which phenocopies a single *bfmR* deletion [35, 36, 44] and shows reduced capsule levels compared to WT and the hyper-encapsulated *bfmS*\* mutant (Fig 5B). As predicted by Tn-seq, Δ*bfmRS* is impervious to Loki in both liquid challenge and plaque assays (Fig 5C, D). Given this non-susceptibility and the epistatic relationship of *bfmR* to *bfmS*, we predicted that spontaneous phage-resistance mutations in the hypersusceptible *bfmS*\* strain would map to *bfmR*. To test this prediction, mutants of *bfmS*\* forming colonies within zones of lysis by Loki were isolated and analyzed by whole-genome sequencing. In two mutants, the resistance mutations were indeed in *bfmR*, causing amino acid substitutions G100D and T85I. Each mutation completely prevented Loki infection in either liquid (Fig 5C) or plaque (Fig 5D) assays, despite the hypersusceptibility of the parent strain. G100 and T85 are highly conserved residues near the BfmR dimerization interface and the predicted site of phosphorylation, respectively, based on the published crystal structure [57, 58]. T85 also aligns with the “switch” residue of *E. coli* CheY (T87), in which a similar mutation to isoleucine blocks phosphorylation-induced conformational changes and activation [59]. G100D and T85I are thus predicted to inactivate BfmR. Consistent with inactivation, both substitutions greatly reduced the phosphorylated (predicted active [44]) fraction of the protein (BfmR∼P, Fig 5E) as well as total BfmR amounts (S5A Fig), as analyzed in phosphate affinity shift gels (Materials and Methods). The BfmR substitutions also reduced capsule synthesis to levels seen with Δ*bfmRS* (Fig 5B). We note that in other Loki-escape derivatives of *bfmS*\*, *gtr6* was again a site of mutations, including two independent 440A_9→8_ mutations and another, novel slipped-strand mispairing mutation within a different poly-A tract (384A_8→7_) (Fig 3D, S2 Table).

**Fig. 5.**
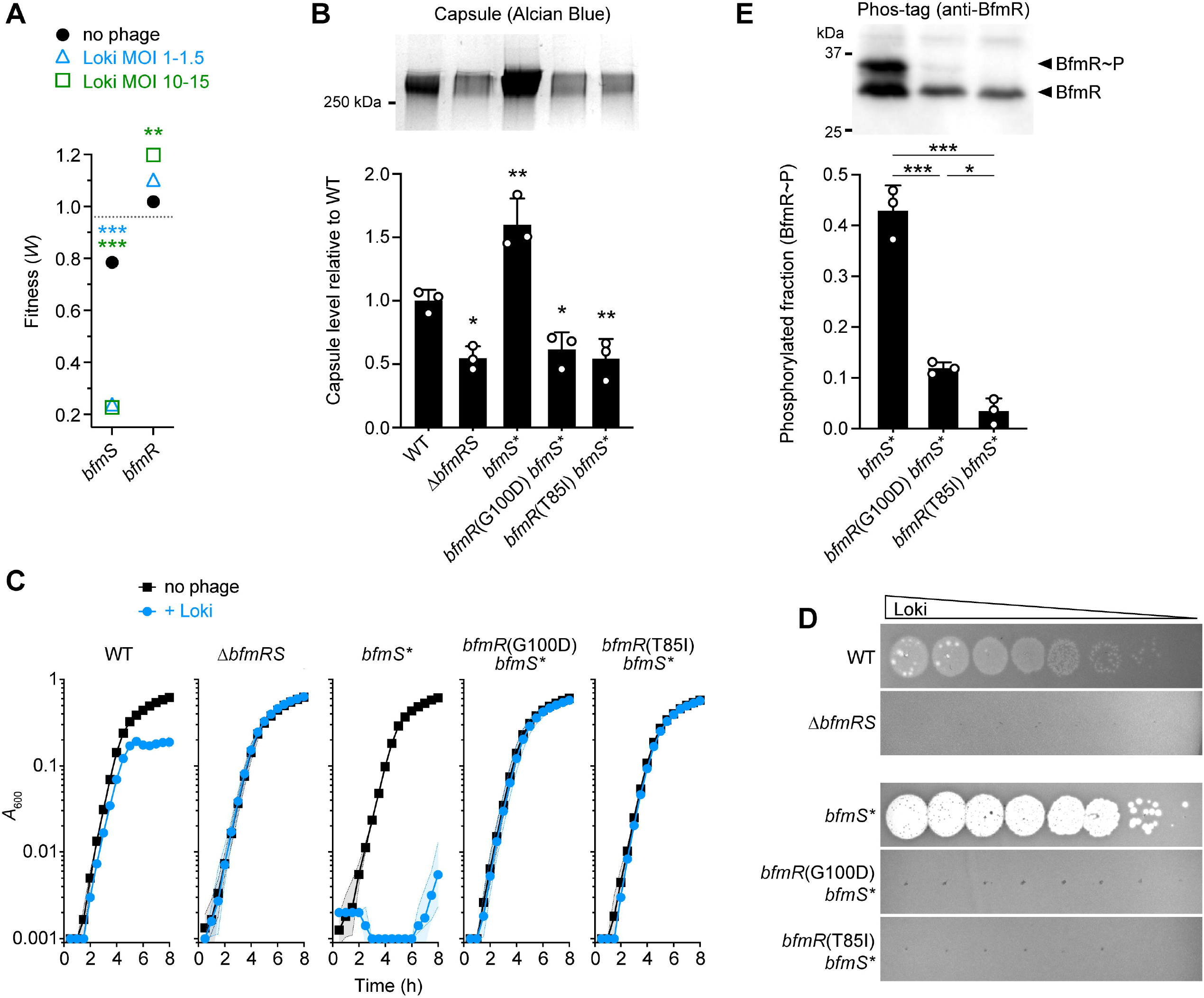
Susceptibility to Loki depends on BfmRS activity and correlates with control of capsule levels. (A) *bfmS* and *bfmR* transposon mutations have opposing effects on Tn-seq fitness during Loki challenge. Data points show average gene-level Tn-seq fitness values (*W*) with Loki infection at indicated MOI vs uninfected control, analyzed by unpaired t-tests. Dotted line indicates the genome-wide average *W* during Loki challenge (MOI 10-15). (B) Capsule production by different *bfmRS* mutants. Bars show mean capsule level ± s.d. (n=3), analyzed by one-way ANOVA with Dunnett’s multiple comparisons test (mutant vs WT). A representative gel is shown above the graph. (C) Liquid challenge and (D) plaque formation assays with Δ*bfmRS* and *bfmR* receiver domain point mutants. Liquid challenge data are presented as in Fig 1A (n=3). (E) BfmR phosphorylation level was analyzed in cell lysates of receiver domain mutants vs the parent control (*bfmS*\*) by Phos-tag western blotting. BfmR∼P as a fraction of total BfmR was quantified from n=3 samples. Bars show mean ± s.d., analyzed by one-way ANOVA with Tukey’s multiple comparisons test. A representative blot is shown above the graph.

The opposing effects on capsule and Loki susceptibility of *bfmR* (low capsule, low infectivity) and *bfmS* (high capsule, high infectivity) mutants are consistent with regulation of the capsule receptor contributing to the differential sensitivity to Loki in each strain. In agreement with this model, compared to WT, Loki showed reduced adsorption to Δ*bfmRS* and dramatically increased adsorption to *bfmS*\* (slow vs rapid decline in the fraction of free phage, respectively, Fig 4B). The increased adsorption and infectivity of Loki conferred by *bfmS* mutation depended on the K3 capsule, based on two lines of evidence. First, deleting *itrA* or *gtr6* in 17978 *bfmS* mutants blocked Loki adsorption (S6A Fig) and killing (S6B Fig). Second, *bfmS* mutation greatly enhanced Loki plaque formation in 19606 (K3), but had no effect on Loki susceptibility in AB5075 (K25) (S6C Fig). These data thereby further support the notion that the BfmRS-dependent boost in Loki susceptibility depends on capsule.

### Defects in RnaA and DsbA control capsule and phage susceptibility through BfmRS

We reasoned that additional lesions uncovered in our genome-wide screen may owe their altered Loki susceptibility to modulation of the capsule receptor. We found that knockouts of *rnaA* and *dsbA* in 17978, which hypersensitize the bacteria to Loki, also each increased capsule production (by ∼50%, Fig 6A) and phage surface adsorption (Fig 4B), similar to the enhancements caused by *bfmS* mutation. As expected, augmented capsule production was reversed by expressing each WT allele in trans (S1B Fig).

**Fig 6.**
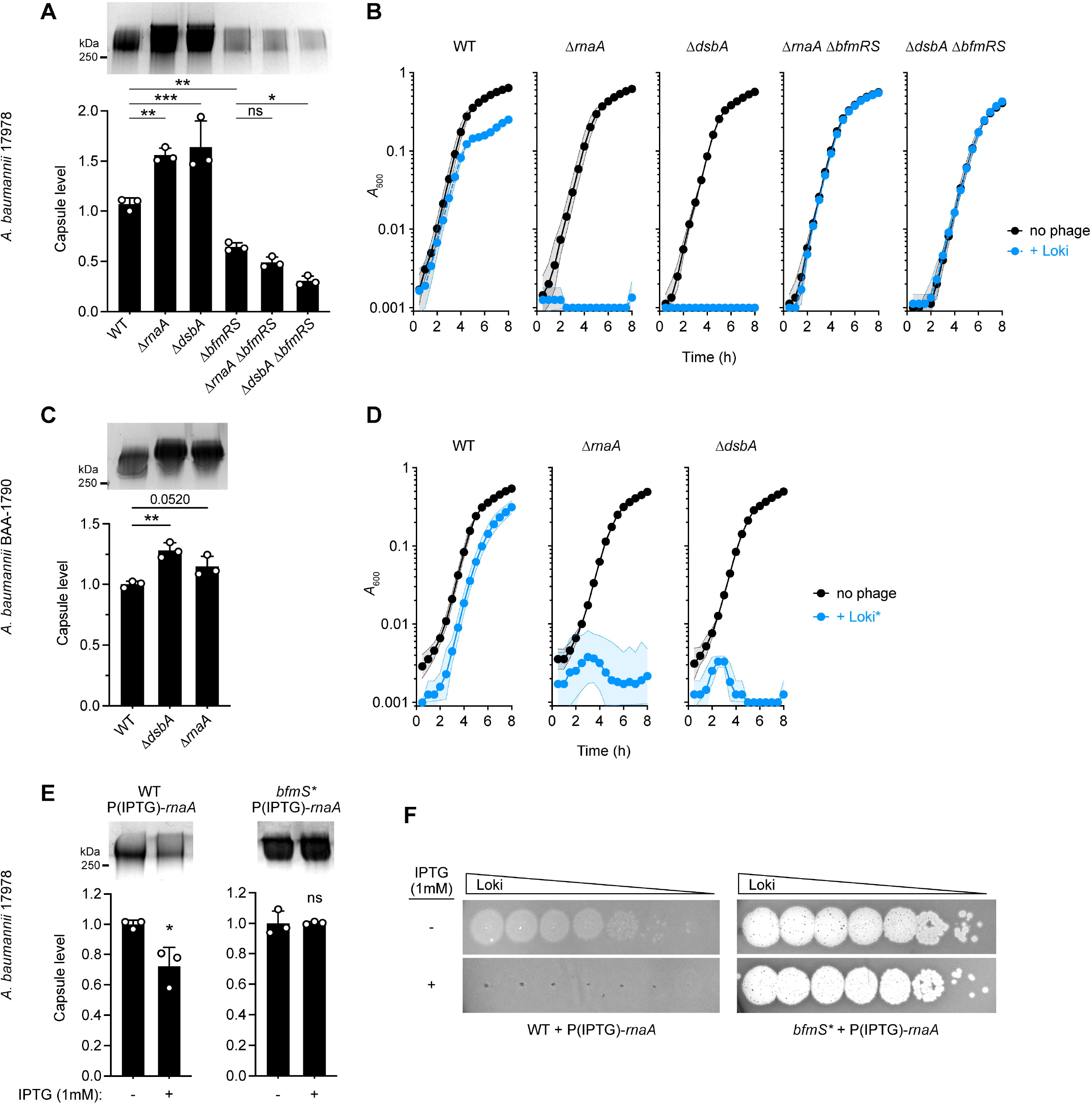
RnaA and DsbA modulate capsule production and Loki susceptibility through BfmRS and in different clinical isolates. (A) Capsule levels measured in 17978 derivatives with Δ*rnaA* and Δ*dsbA* mutation singly and in combination with Δ*bfmRS*. A representative gel (top) and quantification from n=3 samples (bottom; means ± s.d. analyzed by one-way ANOVA with Šídák’s multiple comparisons test) are shown as in Fig 5B. (B) Liquid challenge assays examining susceptibility of strains in A to Loki attack. Data presented as in Fig 1A (n=6). (C-D) Δ*rnaA* and Δ*dsbA* derivatives of carbapenem-resistant *A. baumannii* strain BAA-1790 show increased capsule and Loki sensitivity. Capsule levels (C) were analyzed as in A. Bacterial growth during liquid challenge with a virulent Loki derivative (Loki*) (D) was analyzed as in B. (E-F) *rnaA* overexpression via high-copy plasmid decreases capsule production and blocks Loki infection in a manner dependent on the BfmS signal receptor. WT or *bfmS*\* 17978 bacteria harbored pJE127 (IPTG-dependent *rnaA*) and were grown in the presence or absence of IPTG. Representative capsule gel (E, top), capsule quantification from n=3 samples (E, bottom), and Loki plaque assays (F) are shown. Mean capsule levels ± s.d. were analyzed by unpaired t-tests. P values: *, ≤0.05; **, ≤0.01; ***, <0.001. ns, not significant.

To determine if the link between capsule biosynthesis, phage susceptibility, and *rnaA* and *dsbA* applies to a different K3 strain, we also deleted the genes in BAA-1790. Similar to their effects in 17978, each deletion caused BAA-1790 to hyperproduce capsule (Fig. 6C) and increase susceptibility to Loki (S7A Fig). Because the BAA-1790 WT parent showed complete resistance to Loki in the liquid challenge assay, we also tested a mutant derivative of Loki (Loki*, Materials and Methods) with increased virulence against WT strains (S7B Fig and Fig. 6D, WT). Δ*rnaA* and Δ*dsbA* each caused BAA-1790 to be completely inhibited by Loki* under conditions in which the WT showed minimal inhibition (Fig. 6D). RnaA and DsbA lesions thus amplify capsule and phage susceptibility in both a laboratory strain carrying KL3 as well as a more recent, carbapenem-resistant clinical isolate carrying KL22, suggesting that the proteins act through a conserved envelope modulation pathway.

Given the highly similar capsule and phage susceptibility phenotypes shared by *rnaA*, *dsbA*, and *bfmS* mutants, and the ability of the BfmRS regulatory system activity to control these phenotypes, we considered the possibility that *rnaA* and *dsbA* mutations act through BfmRS. We thus defined the epistatic relationship of the TCS to *rnaA* and *dsbA* in 17978. We found that combining Δ*rnaA* or Δ*dsbA* with Δ*bfmRS* led to low capsule levels (Fig 6A) and complete resistance to phage attack in both liquid challenge (Fig 6B) and plaque formation assays (S1C Fig), mirroring the Δ*bfmRS* single mutant phenotypes. In addition, we examined how *rnaA* overexpression interacts with the TCS, taking advantage of the above observation that overexpression of *rnaA*, but not *dsbA*, not only complements the corresponding deletion but also lowers capsule and Loki susceptibility below WT levels (Fig 2D; S1A,B Fig). We predicted that these decreases would depend on BfmRS. We found that overexpressing *rnaA* in trans in WT led to lower levels of capsule compared to uninduced control (Fig 6E), and completely prevented Loki plaque formation (Fig 6F). Both of these effects were blocked when *rnaA* induction occurred in a *bfmS*\* strain harboring the inactive version of the BfmS receptor (Fig 6E, F). In contrast to the results with *rnaA* and as predicted from Fig 2D, overexpressing *dsbA* in WT did not inhibit plaque formation (S1D Fig). Together, these results are consistent with the model that RnaA and DsbA defects act through BfmRS to modulate capsule and Loki infectivity, with the TCS sensing relative levels of RnaA and presence/absence of DsbA function.

Several additional lines of evidence support the model that BfmRS responds to RnaA and DsbA defects. First, Δ*rnaA* and Δ*dsbA* mutants increased the transcription of capsule biosynthesis genes (*wza* and *gnaA*, Fig 7A) and the activity of a BfmRS-dependent promoter fused to GFP (18040p-GFP, Fig 7B), mirroring the effects of *bfmS* mutation on the same phenotypes [36]. Similar increases were not seen with a BfmRS-independent reporter construct (*adc*p-GFP [36], Fig 7B), or with 18040p-GFP when Δ*rnaA* or Δ*dsbA* were combined with Δ*bfmRS* (Fig 7C). The reporter results indicate that transcriptional enhancement by RnaA and DsbA deficiency is specific and dependent on BfmRS. Second, Δ*rnaA* and Δ*dsbA* each increased the fraction of BfmR∼P by approximately 3-fold (Fig 7D) and appeared to increase total BfmR levels as well (S5B Fig), similar to the effects of *bfmS* deletion. Third, analysis of previous genome-wide fitness datasets [41] revealed that *rnaA*, *dsbA,* and *bfmS* mutants have highly correlated antibiotic susceptibility profiles (phenotypic signatures), which are likely to reflect functional connectivity between the genes (Fig 7E and S1E Fig, r = 0.58-0.67, p < 0.0005). The *bfmS*, *rnaA*, and *dsbA* signatures also positively correlated with those of other *A. baumannii* Dsb system homologs (DsbB, DsbD, and at least one of 3 DsbC paralogs, ACX60_RS16880), suggesting that disulfide bond formation in general is linked to the functions of BfmS and RnaA. Mutations in *dsbB*, *dsbD* and the same *dsbC* paralog (RS16880) also showed decreased Tn-seq fitness during Loki challenge, albeit not to the degree seen with *dsbA* (Fig 7F). Fourth, our previous RNA-seq data [36] showed that expression of several *dsb* genes as well as *rnaA* is under the control of BfmRS, with *bfmR* and *bfmS* mutants having differential effects on their transcription (S1F Fig). Finally, while the Δ*rnaA*, Δ*dsbA*, and Δ*bfmRS* individual mutants and the Δ*rnaA* Δ*bfmRS* combined mutant do not show substantially impaired growth at 37°C, we found that combining Δ*dsbA* and Δ*bfmRS* results in a synthetic growth defect (Fig 7G). This result implies that BfmRS protects *A. baumannii* from DsbA deficiency and the consequent defects in periplasmic protein disulfide bonds, consistent with a toxicity-stress response relationship.

**Fig 7.**
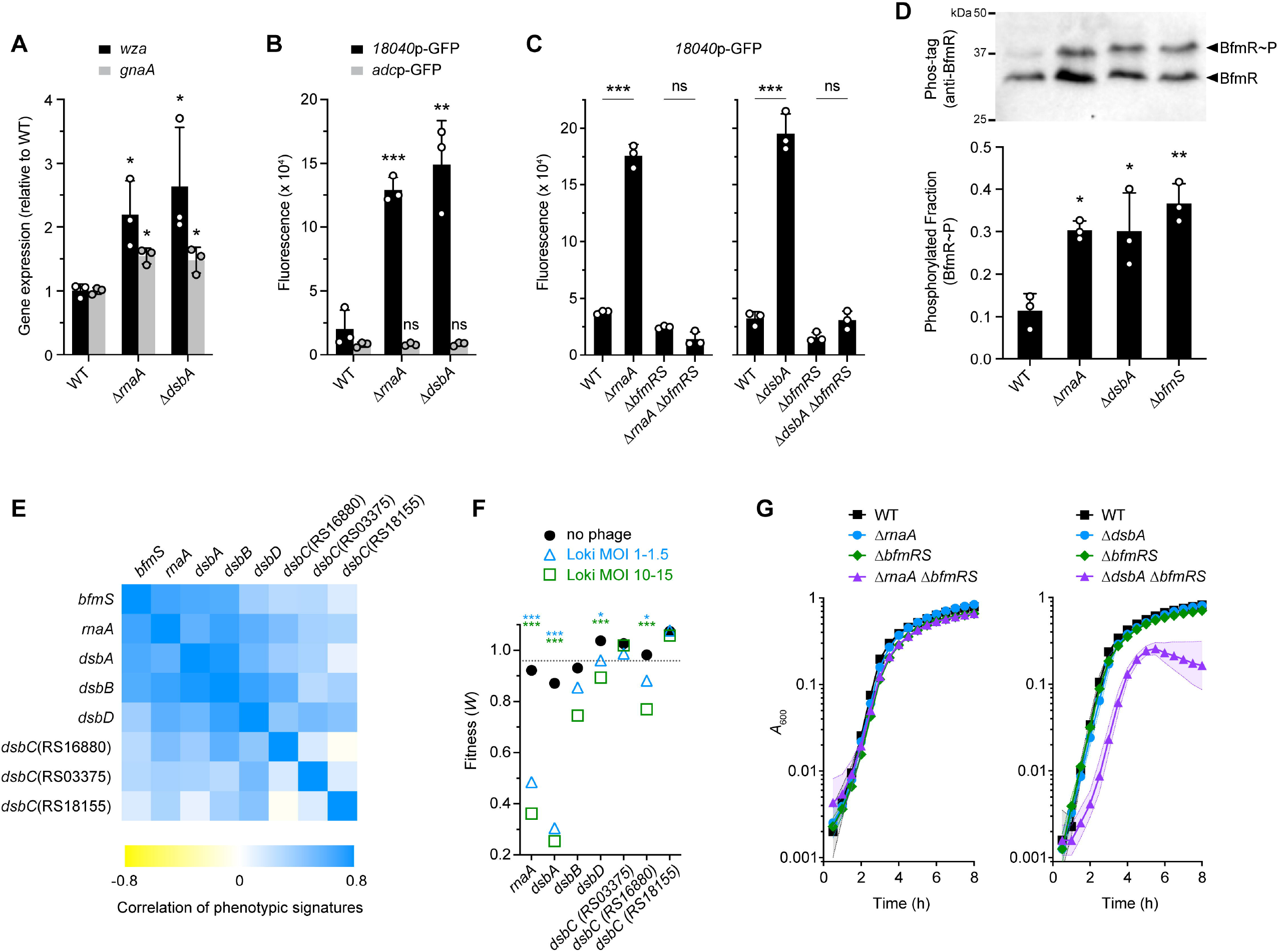
RnaA/DsbA deficiency activates BfmRS. (A) Transcription levels of capsule biosynthesis genes *wza* (black) and *gnaA* (gray) measured by qRT-PCR in Δ*rnaA* and Δ*dsbA* mutants. Bars show mean fold change vs WT ± s.d. (n=3); analyzed by unpaired t-tests comparing mutant vs WT. (B-C) Fluorescence reporter assays. Strains harbored a GFP transcriptional fusion to the noted promoter region and reporter fluorescence was measured as fluorescence units/A_600_. (B) Δ*rnaA* and Δ*dsbA* activation of a BfmRS-dependent (*18040*p, black), but not BfmRS-independent (*adc*p, gray) promoter. (C) Δ*rnaA*/Δ*dsbA* activation of *18040*p requires BfmRS. Bars show mean ± s.d. (n = 3). Means were analyzed by unpaired t test comparing mutant vs WT (B) or by one-way ANOVA with Šídák’s multiple comparisons test (C). (D) Phos-tag analysis of BfmR phosphorylation. Δ*rnaA* and Δ*dsbA* increase BfmR∼P to a level matching that caused by the hyperactive Δ*bfmS* allele. Presented as in Fig 5E (n=3). Asterisks show results of one-way ANOVA with Tukey’s multiple comparisons test (mutant vs WT); comparisons between mutants were not significant. (E) Heat map shows the Pearson correlation coefficients (r) analyzing relatedness of Tn-seq phenotypic signatures [41] of *bfmS, rnaA,* and *dsb* genes. (F) Tn-seq fitness of *rnaA* and *dsb* transposon mutants in the phage challenge screen. Data presented as in Fig 5A. (G) Δ*dsbA* but not Δ*rnaA* shows a synthetic growth defect with Δ*bfmRS.* Bacteria were incubated in LB at 37°C and growth measured as in Fig 1A (n=3). P values: *, ≤0.05; **, ≤0.01; ***, <0.001. ns, not significant.

Sequence analysis of RnaA, an RNase-domain protein, sheds light on its connection to DsbA and BfmRS. RnaA contains a signal peptide as well as eight cysteine residues (S8 Fig). These features potentially place RnaA in the periplasm where internal disulfide bonds could be formed by DsbA, as occurs with its *E. coli* ortholog, RNase I [60]. Unlike RNase I, however, RnaA and its close orthologs within *Acinetobacter* and *Psychrobacter* completely lack the catalytic histidines and other key residues in its two conserved active sites (CAS) critical for RNase activity (S8 Fig) [61], as previously noted with the *A. baumannii* RnaA sequence [62]. These findings suggest that RnaA may be a pseudoenzyme, whose levels and/or folding may influence BfmS signaling and phage susceptibility.

### Factors influencing phage infection independent of capsule modulation

While modulation of capsule is a major feature of altered Loki susceptibility, we found that other hypersusceptibility determinants—including *gtrOC3*, *gtrOC4*, and *lon*—affect vulnerability to Loki independently of regulating the capsule receptor. Deletions of these genes resulted in growth defects during phage challenge in liquid medium that were characterized by different kinetics compared to those of the hypo-encapsulated *rnaA*, *dsbA*, and *bfmS* mutants (Fig 2B, C), consistent with an alternate mechanism underlying their susceptibility defects.

Deletions of *gtrOC3* and *gtrOC4*, which encode predicted LOS outer core glycosyltransferases [29], resulted in the expected enrichment of truncated forms of LOS without lowering the overall amount of LOS produced (Fig 8A). The deletions also showed no change to the levels or size distribution of capsular polysaccharide compared to the WT control (S4B Fig). Despite having WT levels of the capsule receptor, the Δ*gtrOC3* and Δ*gtrOC4* mutants both showed similar enhancements to Loki adsorption, particularly at the later 10 and 20 minute time points (Fig 4B). As noted above, mutating other LOS outer core synthesis genes also decreased fitness during Loki challenge compared to controls, albeit to a lesser extent than *gtrOC3* and *gtrOC4* (Fig 8B, S1 Dataset). This suggests that a range of outer core defects can affect Loki susceptibility. By combining additional mutations with Δ*gtrOC3*, we found that its enhanced Loki attachment and susceptibility phenotypes still depended on initial binding to capsule. A Δ*gtrOC3* Δ*itrA* double mutant showed complete lack of Loki adsorption (S6D Fig), and Δ*gtrOC3* Δ*itrA* and Δ*gtrOC3* Δ*gtr6* double mutants showed full phage resistance in solid and liquid media (Fig 8C, D). The full-length LOS outer core thus functions to inhibit the ability of Loki to efficiently adhere to and infect *A. baumannii*, likely after the initial requisite interaction with capsule.

**Fig 8.**
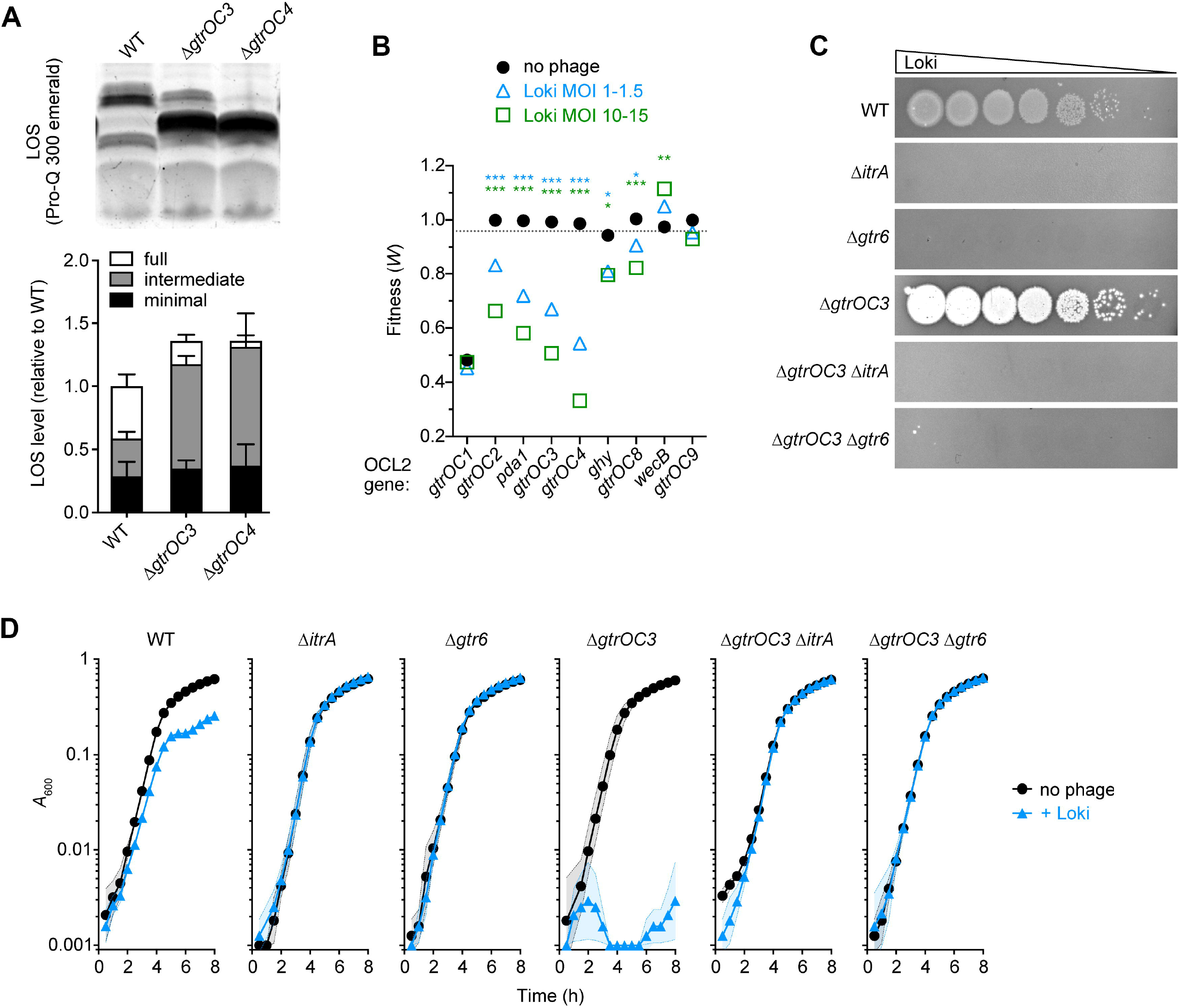
The full-length LOS outer core impedes attack by Loki independent of changes to capsule. (A) Δ*gtrOC3* and Δ*gtrOC4* cause enrichment of a truncated LOS intermediate. A representative LOS gel (top) and quantification from triplicate samples (bottom) are shown. Bars indicate mean normalized LOS levels ± s.d.; one-way ANOVA showed no significant difference in total LOS levels among strains (P=0.34). (B) Tn-seq fitness of LOS outer core synthesis locus (OCL2) mutants in the phage challenge screen. Data presented as in Fig 3A. (C-D) Enhancement of Loki susceptibility in *gtrOC* mutants depends on capsule. (C) Plaque formation assay with Δ*gtrOC3,* Δ*itrA*, Δ*gtr6*, and double mutants. (D) Liquid challenge assay with strains in C (n = 3). P values: *, ≤0.05; **, ≤0.01; ***, <0.001. ns, not significant.

Deleting *lon* caused no significant change to capsular polysaccharide levels (S4C Fig), consistent with previous observations [63], and had no effect on LOS quantity or banding pattern (S9C Fig). As predicted from the lack of major changes to the key surface components, Loki adsorption was not significantly enhanced with the Δ*lon* mutant vs WT; rather, adsorption appeared lowered at the early 5-min time point compared to WT (Fig 4C). Lon protease may thus be affecting susceptibility to Loki by influencing a lytic cycle step occurring after surface interactions.

We considered the possibility that the altered Loki susceptibility of mutants showing altered capsule production could also involve alterations to LOS. We therefore examined LOS from the *bfmRS*, *bfmS*, *rnaA* and *dsbA* deletion mutants. LOS from Δ*bfmRS*, Δ*rnaA* and Δ*dsbA* showed no differences in banding pattern or total levels compared to LOS from WT (S9A,B Fig). While overall amounts of LOS from Δ*bfmS* were not significantly altered, we observed the appearance of a single, novel band migrating slightly faster than the full-length LOS (S9A Fig, arrowhead). This band may represent an LOS intermediate, although it is slower migrating and much less abundant than the major intermediate band enriched in Δ*gtrOC3* and Δ*gtrOC4* mutants. Thus, while altered capsule receptor abundance is a key feature of the enhanced Loki susceptibility of BfmRS-hyperactive bacteria, a contribution of altered LOS to the dramatic hypersusceptibility of *bfmS* mutants cannot yet be ruled out.

### Enhanced phage multiplication caused by BfmRS hyperactivity and Lon deficiency

To examine how phage hypersusceptibility loci affect Loki intracellular replication, we performed one-step phage growth analysis (Materials and Methods). The results of these experiments are shown in Table 1 and S10 Fig. Infection of WT bacteria resulted in a burst size of ∼20 plaque-forming units (PFU) per infective center and an eclipse period of ∼40 minutes. Compared to this control, *bfmS*\* led to a dramatic amplification in phage replication, increasing burst size >10-fold and shortening the eclipse by half. BfmRS-hyperactive Δ*rnaA* and Δ*dsbA* mutations had similar effects, increasing the burst 3-5-fold and halving the eclipse period. These results indicate that the BfmRS pathway shapes phage susceptibility not only by affecting adsorption but also by controlling intracellular phage replication. Δ*lon* also increased Loki burst size (by 7-fold) and shortened the eclipse phase by ∼10min, consistent with the above idea that Lon primarily influences post-adsorption phage growth. By contrast, Δ*gtrOC3* showed no change in the magnitude of phage replication, indicating that LOS outer core synthesis mainly affects phage adsorption.

**Table 1.** Results from one-step growth experiments.

| Host strain<sup>a</sup> | eclipse period (min)<sup>b</sup> | Burst size |  |  | n |
| --- | --- | --- | --- | --- | --- |
|  |  | mean ± SEM (PFU) |  | adjusted p-value<sup>c</sup> |  |
| WT | 40 | 20.6 | ± 1.2 |  | 12 |
| *bfmS* * | 20 | 225.9 | ± 17.9 | <0.0001 | 4 |
| *ΔrnaA* | 20 | 61.8 | ± 9.9 | 0.0069 | 4 |
| *ΔdsbA* | 20 | 94.9 | ± 6.0 | <0.0001 | 4 |
| *ΔgtrOC3* | 30 | 27.3 | ± 1.8 | 0.9843 | 4 |
| *Δlon* | 30 | 141.2 | ± 19.7 | <0.0001 | 4 |

a. *A. baumannii* 17978 background
b. mode value from experiments using 10-minute sampling intervals
c. from one-way ANOVA with Šídák's multiple comparisons test (comparing each mutant vs WT)

## Discussion

Bacteriophage show promise as an alternative therapy against drug-resistant bacteria, but how vulnerability to phage attack can be controlled is poorly understood. In this work we have defined the genome-wide mechanisms of *A. baumannii* susceptibility to killing by phage Loki. Our results support a model in which this susceptibility is determined by numerous major factors affecting the efficiency of phage adsorption and post-adsorption replication (Fig. 9A). We found that capsule type K3 provides the primary Loki receptor. LOS outer core glycans antagonize adsorption likely by impeding access to a secondary binding site, which could be membrane-proximal residues of the capsule or of the LOS molecule (e.g., the conserved LOS inner core). Phage adsorption as well as intracellular phage replication are controlled in dual fashion by the BfmRS regulatory system (Fig. 9A). Activating the TCS augments capsule production and phage growth, while deactivating the TCS lowers capsule and confers complete resistance (Fig. 9B, C). Lon is another defense factor that specifically limits phage multiplication (Fig. 9A, C). How BfmRS and Lon control phage growth dynamics intracellularly remains to be defined. Of note, the hypersusceptibility seen with *A. baumannii* mutants showing capsule overproduction or Lon deficiency stand in contrast to the situation in *E. coli,* in which these features are associated with broad phage resistance [64, 65]. Finally, alteration of capsule structure through naturally occurring glycosyltransferase mutations provides an additional pathway to phage resistance through loss of adsorption (Fig. 9B). The *gtr6* locus appears to be a hotspot for such mutations, related to strand slippage and insertion sequence elements, that could potentially facilitate variation of capsule structure within bacterial populations [66, 67].

**Fig. 9.**
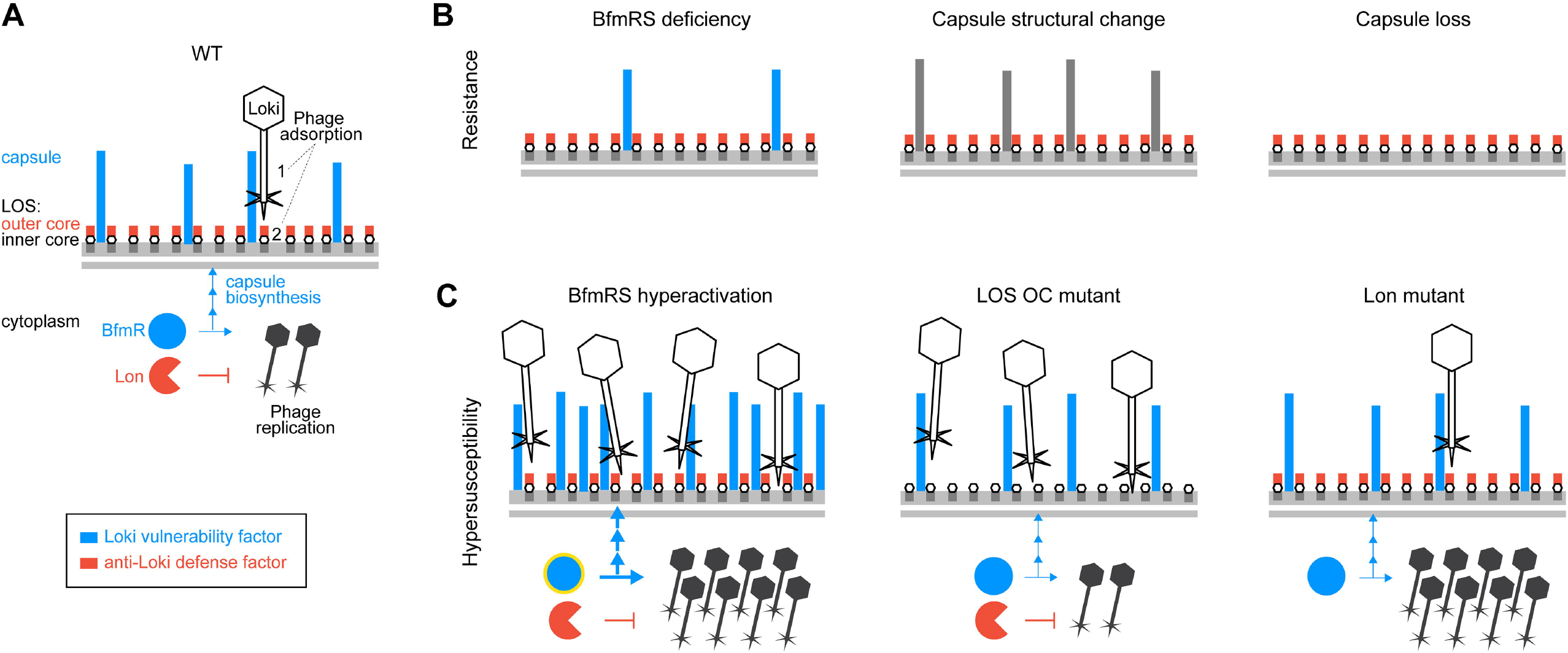
Model for modulation of phage infection by multiple susceptibility factors in *A. baumannii*. Vulnerability factors that act to enhance Loki infection are shown in blue; defense factors that antagonize Loki infection are shown in red. Additional candidate defense factors, including the methyltransferase AamA or transcription factor DksA, were identified through screening but require confirmation and are not shown. (A) Loki infection of WT bacteria. Loki adsorbs in two steps: (1) interaction with the primary adsorption receptor, capsular polysaccharide of type K3, and (2) interaction with an LOS-associated secondary site. This site may be the conserved LOS inner core, or LOS-proximal capsule residues, and access is antagonized by the LOS outer core. BfmRS enhances Loki infection by positively regulating both the capsule receptor (via transcriptional activation of capsule biosynthesis) and intracellular phage replication (by unknown mechanisms). Lon protease negatively regulates intracellular phage replication, by unknown mechanisms. (B) Loki resistance arises from capsule receptor down-regulation (BfmRS deficiency), structural alteration (via mutation of hotspot gene *gtr6*), or complete loss (mutation of a core biosynthesis gene, e.g. *itrA* or *wzx*), blocking phage adsorption. BfmRS deficiency may also reduce phage intracellular replication. (C) By contrast, Loki hypersusceptibility arises from hyperactivity of vulnerability factors or loss of defense factors. Hyperactivation of BfmRS, defined by increased BfmR∼P (yellow halo), results from *bfmS*, *dsbA*, or *rnaA* mutations and may also be triggered by external stresses related to oxidative protein folding. BfmRS hyperactivity increases the level of the capsule receptor, enhancing phage adsorption, while also stimulating phage intracellular multiplication. Loss of LOS OC sugars increases binding of Loki to surface sites without altering its intracellular multiplication. Loss of Lon increases phage multiplication without increasing adsorption.

*A. baumannii* phages frequently target specific host capsules [26, 35–38]. That capsule structural modulation mediates phage escape has implications for strategies to control the evolution of *A. baumannii* phage resistance and virulence. Mutations in *gtr6*, which determines the addition of N-acetyl-β-D-glucosamine branches on the K3 capsule backbone [26], are known to amplify virulence [26]. Our findings thus suggest that phage, in the environment or during therapy, may impose selective pressures facilitating the evolution of altered capsules with the potential collateral effect of increasing pathogenicity. Notably, a repeat region within another predicted side-chain glycosyltransferase, *gtr76*, was a site of mutations in phage-resistant K116 *A. baumannii* isolated from a patient receiving phage therapy [23], although effects on capsule structure were not determined. Mutational capsule loss is an additional, well-described phage escape mechanism (Fig. 9B) usually associated with lowered *in vivo* fitness and other trade-offs, making this form of resistance an appealing target of sequential or combination therapies exploiting the trade-offs [18, 68]. Our data, and potentially those of [23], argue that capsule-loss mutations should not be considered a universally predictable path of *A. baumannii* phage resistance and that capsule-altered mutants maintaining (and possibly enhancing) virulence should also be considered. This may be particularly relevant when targeting specific capsules such as K3. Combining phages to target both Gtr6^+^ and Gtr6^-^ [56] capsules may allow treatment of K3 *A. baumannii* infections while blocking the rise of undesired mutants. In sum, these observations support the notion that prior research into resistance mechanisms can enhance the predictability of bacterial evolution during phage exposure and may allow improvements to the therapeutic uses of phage [18, 23, 68].

Controlling the abundance of capsule is also tightly connected to modulation of virulence, and we show here that this control has the additional effect of modulating phage susceptibility, with implications for developing phage-drug synergies [68–70] and for understanding variable capsule production in the pathogen. *A. baumannii* regulates capsule synthesis through multiple mechanisms, including the BfmRS stress response and stochastic phenotypic switching systems, and the regulatory states associated with higher capsule production show increased virulence [35–38]. We show that BfmRS hyperactivity is an Achilles’ heel that can be exploited by capsule-targeting phages, allowing them to bind, multiply, and kill with enhanced efficiency. It is possible that antimicrobials that increase capsule [35] could similarly potentiate phage activity, while the opposite may be seen with antagonists of BfmRS/capsule production. Chemical inhibitors of DsbA [46, 71], which would activate BfmRS and capsule synthesis while impairing envelope proteins generally, represent an attractive candidate strategy for small molecule potentiation of phage attack. Regarding the basis of variable capsule production, our results raise the possibility that environmental phage are among the putative selective pressures that enrich low-capsule (translucent) variants arising from stochastic phenotype switching [37].

By examining determinants of Loki susceptibility acting through BfmRS, this work provides new leads into the signals detected by the virulence-resistance regulator, which have remained poorly defined. We reveal that changes in DsbA and RnaA generate signals sensed by the TCS, leading to transcriptional control of capsule and phage infectivity. The finding that lesions affecting periplasmic disulfide bonds generate a stress-response has a parallel in the *E. coli* Cpx system [72]. *dsbA* mutants activate Cpx [73], which in turn regulates *dsbA* in a feedback loop [74, 75]. In *A. baumannii*, expression of *dsb* and oxidative stress response genes is modulated by BfmRS [36, 44]. These findings reinforce the view that BfmRS has a subset of activities analogous to those of Cpx [24]. As noted above, several observations connect DsbA to RnaA. A major substrate of DsbA in *E. coli* is the RnaA ortholog RNase I, a non-essential, periplasmic, broad-substrate endonuclease [60]. RNase I and RnaA both have eight cysteines, but RnaA is likely non-catalytic, and its levels serve as a BfmRS signal. A possible interpretation is that *Acinetobacter* has coopted its RNase T2 family protein to function as a sensor of envelope disulfide bond defects and/or oxidative damage. It is worth noting that a sensor of disulfide bond defects has been reported in *E. coli*, NlpE, which transmits signals to Cpx [73]. *A. baumannii* has an NlpE ortholog but it lacks the domain implicated in disulfide sensing [76]. This suggests that the pathogen uses a distinct strategy to monitor periplasmic disulfides, with consequences for vulnerability to phage. Future work will focus on investigating this strategy and defining the connections between RnaA, oxidative folding stress, and BfmS in protecting and altering the *A. baumannii* cell envelope.

In addition to capsule control pathways, our data with LOS outer core glycosyltransferase mutants support an important role for outer core glycans in defending against Loki attack. It is possible that the outer core hinders phage binding to a more internal secondary receptor. Another possibility is that Loki’s preferred secondary receptor is a truncated form of LOS, whose levels increase in *gtrOC* mutants, increasing adsorption efficiency. Previous work showed that mutation of *lpxA*, which completely prevents LOS production, is also reported to block Loki adsorption [40]. This could be explained by the loss of a putative secondary binding site as noted above, and/or by loss of capsule when LOS synthesis is completely arrested [77]. Examination of additional LOS variants should assist in defining the precise mechanisms by which LOS participates in phage infection and defense.

In conclusion, we have analyzed the genome-wide determinants of susceptibility to bacteriophage Loki in *A. baumannii* and identified multiple mechanisms that modulate phage infectivity, including several linked to altered bacterial virulence. Our results expose a vulnerability in a pathogenic strategy of *A. baumannii*, capsule hyperproduction, that allows enhanced killing by the capsule-specific phage. Additional work is needed to define the novel stress sensing mechanisms that impact on phage virulence, how cytoplasmic factors control intracellular viral multiplication, and how the susceptibility factors identified here affect infection by additional families of *Acinetobacter* phages. Future studies focusing on manipulation of phage hypersusceptibility pathways in *A. baumannii* may facilitate the development of improved phage-based combination therapies targeting the pathogen.

## Materials and Methods

### Strains and growth conditions

Bacterial strains used in this work are described in S1 Table. *A. baumannii* strains were derivatives of ATCC 17978 unless otherwise noted. Bacteria were cultured in Lysogeny Broth (10 g/L tryptone, 5 g/L yeast extract, 10 g/L NaCl) (LB). Cultures were incubated at 37°C in all cases except phage infection experiments, which used 30°C unless otherwise indicated. LB + 5mM CaCl_2_ was used as culture medium in all phage infection experiments. Liquid cultures were aerated with orbital shaking or with rotation via roller drum, with growth monitored by measuring optical density (OD) as absorbance at 600nm (*A*_600_). LB agar (1.5% w/v) plates were supplemented with antibiotics as needed (tetracycline [Tc] at 10 μg/ml; carbenicillin [Cb] at 50-100 μg/ml, kanamycin [Km] at 10-20 μg/ml, gentamicin [Gm] at 10 µg/ml) or sucrose (10% w/v) (Sigma). Isopropyl β-D-thiogalactoside (IPTG) (Sigma) was added to cultures at 0.1-1 mM for induction of P(IPTG)-controlled constructs. Super optimal broth with catabolite repression (SOC) was used in Tn-seq library preparation. Bacteriophage Loki (vB_AbaS_Loki)[40] was a gift from Dann Turner. A derivative of Loki (Loki*) was isolated by purifying phage from a clear plaque forming within a zone of turbid 17978 lysis.

### Molecular cloning and strain construction

Oligonucleotide primers and plasmids used in this study are listed in Table S1. Most sequences were PCR-amplified from the *A. baumannii* 17978 genome and cloned in pUC18 before subcloning to destination plasmids. Clones were verified by DNA sequencing (Genewiz/Azenta). Gene deletions were constructed in-frame by allele exchange with cloned ∼1kb homology arms using conditionally replicating vectors pSR47S or pJB4648 and sucrose counterselection as described [35]. Deletion mutants were verified by verified by colony PCR. An AB5075 *bfmS*::Tn mutant was obtained from the Manoil lab Three-Allele Transposon Mutant Library and verified by screening for TcR and whole-genome sequencing. For complementation and overexpression experiments, sequences were subcloned downstream of the *lacI*^q^-T5*lac*P module [P(IPTG)] on plasmid pYDE152. Plasmids were delivered into *A. baumannii* by electroporation.

### Phage-challenge Tn-seq

Stocks of *A. baumannii* ATCC 17978 *Mariner* transposon mutants constructed previously [43] were thawed, revived for 1 hour in LB, and then back diluted to OD=0.005 (∼2.5 × 10^6^ CFU/ml) in LB with 5mM CaCl_2_ containing Loki at MOI 1 (or 1.5), at MOI of 10 (or 15), or no phage (uninfected control). Cultures were incubated at 37°C until each reached OD ∼1 (about 8 doublings). This took approximately 2 hours for control cultures, with additional time, usually 15-20 minutes, given to the phage treatment arms as needed to reach the target OD. 10 banks of ∼20,000 mutants per bank were analyzed independently. Genomic DNA (gDNA) was extracted (Qiagen DNeasy) from samples before and after the 8 doublings, quantified by a SYBR green I (Invitrogen) microtiter assay, and tagmented (Illumina Nextera). Transposon-adjacent DNA was then amplified to form Illumina libraries as described [41, 42]. Libraries were size-selected (Pippin HT, 275bp-600bp) and sequenced (single-end 50 bp) using primer mar512 [41] on a HiSeq 2500 with high output V4 chemistry at the Tufts University Genomics Core Facility (TUCF-Genomics).

Sequencing read data were processed, mapped to the 17978 chromosome (NZ_CP012004) and pAB3 (NZ_CP012005), and used to calculate individual transposon mutant fitness (*W*) based on mutant vs population-wide expansion in each condition using our published pipeline and formula [41]. Fitness values were standardized across conditions by normalizing to the average value of insertions in neutral sites (pseudogenes and endogenous transposon-related genes) within the genome predicted to have no effects on growth. Gene-level average fitness was calculated by averaging the fitness values of all individual mutants in a given condition across all pools having a transposon in the first 90% of the gene. Difference in gene-level fitness between phage-challenge and control conditions was deemed significant if it had a magnitude >10%, derived from n ≥ 3 individual mutants, and had q-value <0.05 in unpaired, two-tailed t test with FDR controlled by the Benjamini, Krieger, and Yekutieli two-stage step-up method (Prism 8). Fitness values at genomic loci were visualized with Integrative Genomics Viewer [78] after aggregation across all transposon mutant libraries via the SingleFitness script [79].

### Phage susceptibility analysis

i. *Liquid challenge.* Bacterial cultures were diluted to OD=0.003 in LB + 5mM CaCl_2_ and combined with phage at an MOI=1 or with no phage (uninfected control). Bacterial growth was monitored during incubation at 30°C in microtiter plates in an Epoch 2 or Synergy H1M plate reader (Biotek).
ii. *Plaque formation assay.* Bacterial cultures were diluted to OD=1 in 100μl of LB, combined with 5ml of top agar (LB with 5mM CaCl_2_ and 0.4% w/v agar), and poured onto LB agar plates. 10-fold serial dilutions of phage in Saline-magnesium diluent plus gelatin (SMG) buffer (50mM Tris-HCl, PH=7.5, 100mM NaCl, 8mM MgSO4, 0.01% (W/V) gelatin) [80] were spotted onto the top agar, and plates were incubated at 30°C for 16 hours. Plaques were imaged with white light transillumination using a Chemidoc MP (Bio-Rad).
iii. *Phage adsorption.* Host bacteria were cultured in LB + 5mM CaCl_2_ at 30°C to OD=0.1. Loki was added at MOI=0.1 followed by incubation at 30°C. At regular time intervals, 100μl were sampled and mixed with 100μl SMG buffer saturated with chloroform. After centrifugation (8000 x g, 5 minutes), unadsorbed phage within supernatants was enumerated by plaque formation assay. The fraction of free phage was calculated as the number of PFU at each time point divided by the total input PFU.
iv. *One-step phage growth*. Host bacteria were cultured as in *iii* to OD=0.1. Loki was added at MOI=0.05 in 1 ml volume and allowed to adsorb for 5 minutes. Mixtures were centrifuged (4000 x g, 2 minutes) and supernatants containing unadsorbed phage were removed and used to determine unadsorbed PFUs. Pellets were resuspended in sterile LB + 5mM CaCl_2_ and incubated at 30°C. 100µl samples were taken immediately (T0) and at 10-minute intervals over 90 minutes. Samples were combined with 50µl chloroform and used to determine post-adsorption PFUs. Burst size was calculated by dividing the average PFU from samples at the plateau of phage growth by the infected cell number. Infected cell number was calculated by subtracting the unadsorbed PFU from the total input PFU. Data were combined from two independent experiments, each experiment performed in biological duplicate and with a WT host control arm.

### Whole-genome sequencing of spontaneous phage-nonsusceptibility mutants and clinical isolates

Bacterial mutants (derived from *A. baumannii* 17978 WT, *bfmS*\*, or Δ*bfmS*) forming colonies within areas of Loki lysis/plaquing were purified on LB agar. gDNA was extracted and quantified as above. Illumina libraries were prepared by a modified Nextera method [36] and sequenced (single-end 100) on a HiSeq 2500 at TUCF-Genomics. Variants were identified from sequencing reads by Breseq as described [81, 82]. Mutations were verified by PCR amplification and Sanger sequencing (Genewiz/Azenta). EGA10 and EGA65 genomes were sequenced and hybrid assembled at SeqCenter using sequencing data from Illumina (NextSeq 2000, 2×151bp) and Oxford Nanopore platforms. The genome assemblies were used for species confirmation and MLST strain type determination [83]. K and outer core locus types were determined from genome sequence data using bioinformatics tools [84, 85].

### Capsular polysaccharide analysis

Capsular polysaccharide was extracted from processed cell lysates and analyzed by alcian blue following described protocols [35]. Samples were collected from early post-exponential phase cultures (OD 2-3) and centrifuged. Cell pellets were resuspended with lysis buffer [60mM Tris, pH 8; 10mM MgCl_2_; 50μM CaCl_2_; 20μl/ml DNase (NEB) and RNase cocktail (Amersham); and 3mg/ml lysozyme (Sigma)] and incubated at 37°C for 1 hour. Samples were further processed by 3 liquid nitrogen/37°C freeze-thaw cycles and stepwise treatments with additional DNase and RNase (37°C, 30 minutes), 0.5% SDS (37°C, 30 minutes), and Proteinase K (NEB, 60°C, 1 hour). Samples were combined with SDS-PAGE loading buffer, boiled for 5 minutes, and separated on 4-20% Bio-Rad TGX Tris-glycine gels. Capsule was stained with Alcian blue (Sigma), imaged via white light transillumination (ChemiDoc MP), and quantified using Image Lab (Bio-Rad).

### LOS analysis

LOS was analyzed via Tricine gel electrophoresis of cell lysates as described [41]. Log phase bacteria were washed with PBS, resuspended with Novex tricine SDS sample buffer (Invitrogen, 50μl 1X buffer per 0.5 OD units of bacteria), and boiled for 15 minutes. Samples were separated on Novex 16% tricine gels (Invitrogen) and stained with Pro-Q Emerald 300 (Invitrogen). Gels were imaged using UV transillumination (ChemiDoc MP). Total protein detected with subsequent Coomassie Brilliant Blue staining was used for sample normalization. Relative values were calculated by dividing each normalized LOS value by the total normalized LOS levels in WT [41].

### Gene expression analysis

For RT-qPCR, RNA was extracted from log phase bacteria (Qiagen RNeasy), DNase treated (Turbo DNA-free kit, Invitrogen), and reverse transcribed (Superscript II Reverse Transcriptase, Invitrogen). cDNA served as template in qPCR with gene-specific primers (S1 Table) and SYBR Green Master Mix (Applied Biosystems) using a StepOnePlus system according to the manufacturer’s instructions. The amplification efficiency for each pair was tested by standard curves using serial diluted cDNA, and found to be > 97% in each case. Fold change in gene expression was calculated by using the 2^-ΔΔCt^ method with *rpoC* as endogenous control [36]. qPCR was performed in technical duplicate from biological triplicate samples. Samples lacking reverse transcriptase were used to confirm absence of signal from residual gDNA. For fluorescence reporter analysis, bacteria were harvested from OD=1 cultures, resuspended in PBS, and transferred to a black, clear bottom microplate (Corning 3631). Fluorescence intensity (excitation at 480 nm/emission at 520 nm) and *A*_600_ were measured with a Synergy H1M (Biotek). Density-normalized fluorescence units were calculated as fluorescence intensity/*A*_600_.

### Analysis of BfmR phosphorylation

Phosphate affinity electrophoresis was performed following published protocols [86]. 1 OD of mid-log phase bacteria were resuspended in 55µl of BugBuster reagent with 0.1% (v/v) Lysonase (Novagen). Lysis was assisted by repeated pipetting. 18µl of 4X SDS sample buffer (200mM Tris-HCl pH 6.8, 8% SDS, 0.4% Bromophenol blue, 40% Glycerol) was added, and lysates were electrophoresed in Phos-Tag gels [12% acrylamide, 50µM Phos-tag acrylamide (Wako Chemicals), and 100 μM MnCl_2_] at 30mA. After electrophoresis, gels were incubated for 10 minutes in 1X transfer buffer (50mM Tris, 380mM Glycine, 0.1% SDS, 20% Methanol) with 1mM EDTA to remove manganese, followed by additional 10 minutes in transfer buffer without EDTA. Proteins were transferred to PVDF membranes (100V, 1hr). Total protein was detected in the blots with SYPRO Ruby blot stain (Invitrogen) using the manufacturer’s instructions and a GelDoc EZ (Bio-Rad). Blots were incubated in blocking buffer (5% Milk in TBS-T, 1 hour, room temperature) and probed with rabbit anti-BfmR antiserum (1:30,000 or 1:90,000 in blocking buffer, overnight at 4C) and goat anti-rabbit-HRP secondary antibody (Invitrogen, 1:5,000 in blocking buffer, 1 hour at room temperature). Blots were washed 3×10 minutes in TBS-T before and after incubation with secondary antibody, and washed 10min in TBS before developing. Signals were detected with Western Lightning Plus-ECL (Perkin Elmer) and a ChemiDoc MP, and quantified using Image Lab. Rabbit anti-BfmR antiserum was generated by immunizing a rabbit with purified recombinant BfmR according to standard protocols (Pocono Rabbit Farm and Laboratory).

### Phenotypic signature analysis

Tn-seq phenotypic signatures belonging to *bfmS*, *rnaA*, and *dsb* paralogs in *A. baumannii* were extracted from previous genome-wide datasets [41]. Hierarchical clustering was performed by the average linkage method using Qlucore Omics Explorer (3.7). Pearson correlation analysis was performed using Prism 9.

## Acknowledgements

We thank Dann Turner for the gift of phage Loki and Colin Manoil for the AB5075 *bfmS*::Tn strain. We are grateful to Isabella Bailey, Sarah Hudson, and Wenwen Huo for their technical assistance, and members of the Geisinger lab for helpful discussions.

## Supporting information captions

**S1 Fig. Analysis of phenotypes connected to *rnaA* and DSB genes.** (A) Liquid Loki challenge assays show that overexpression of reintroduced *rnaA* by a high inducer concentration (1mM IPTG) not only reverses the Δ*rnaA* phage susceptibility defect, but causes complete non-susceptibility to the phage. “Chr” indicates the chromosomal mutation; “Plasmid” denotes the gene reintroduced under P(IPTG) control via vector pYDE152; - indicates vector with no reintroduced gene. Bacteria were cultured with or without Loki (MOI 1) and IPTG (1mM) as indicated. Data are presented as in Fig. 1A (n=3). (B) Complementation analysis showing reversibility of capsule deficiency in Δ*rnaA* and Δ*dsbA* strains. Capsular polysaccharide levels were analyzed by SDS-PAGE/Alcian Blue after growth with the noted amount of IPTG to induce the reintroduced gene. (C-D) Plaque formation assays. Loki was spotted onto bottom agar containing WT bacteria with the indicated genotype. In D, bacteria harbored pJE101 (IPTG-dependent *dsbA*), and plates contained 0 or 1mM IPTG. (E) Cluster analysis of Tn-seq phenotypic signatures of *bfmS*, *rnaA*, and *dsb* genes. Heat map shows normalized Tn-seq fitness in z-score units for transposon mutants in each gene (rows) grown in different conditions (columns) [41]. Dendrogram at left shows relationship between signatures based on hierarchical clustering. (F) Heat map shows change in RNA-seq transcript levels in the indicated BfmRS mutant compared to WT control. Data were extracted from previous datasets [35] and log2 fold change values displayed as heat map using Prism 9. P values: *, ≤0.05; **, ≤0.01; ***, <0.001.

**S2 Fig. Analysis of Δ*slt* phenotypes.** (A) Liquid challenge assay with WT or Δ*slt* bacteria cultured with or without Loki (MOI 1). Data are presented as in Fig. 1A (n=3). (B) Plaque formation assay with Loki spotted onto bottom agar with WT and Δ*slt* bacteria. (C) Measurement of *rnaA* transcript levels in WT vs Δ*slt* via qRT-PCR. Bars show mean fold change vs WT ± s.d. (n = 3); analyzed by unpaired t-test. *, P ≤ 0.05.

**S3 Fig. Reintroduction of *itrA* in Δ*itrA* mutant restores Loki susceptibility in the liquid challenge assay, and degree of susceptibility increases with increasing *itrA* induction.** (A) Bacteria of the noted genotype were cultured with the indicated level of IPTG with or without Loki at initial MOI of 1. Growth was measured as in Fig 1A (n = 3). (B) Plaque formation assay with Loki spotted on 17978 WT and isogenic Δ*gtr6* mutant.

**S4 Fig. *gtr6*, *gtrOC*, and *lon* mutations do not affect production of capsular polysaccharide.** Bars show mean capsule levels ± s.d. (n=3). Lanes from representative gels are shown above each graph. No significant difference among means by one-way ANOVA (A, P = 0.797; B, P = 0.963) or by t test (C, P = 0.531).

**S5 Fig. Analysis of total BfmR levels in mutant strains.** Total BfmR levels were quantified from anti-BfmR Phos-tag western blots by combining the signal from phosphorylated and unphosphorylated forms of BfmR, and normalizing to total protein in the sample as determined by SYPRO Ruby staining. (A) Graph corresponds to the blot shown in Fig. 5E. (B) Graph corresponds to the blot shown in Fig. 7D. Bars show mean capsule level ± s.d. (n=3), analyzed by one-way ANOVA with Dunnett’s multiple comparisons test (mutant vs WT). *, P value ≤ 0.05.

**S6 Fig. *bfmS* and *gtrOC* mutant hypersusceptibility to Loki depends on capsule biosynthesis.** (A, D) Phage adsorption assays showing that enhancement of Loki adsorption by *bfmS* (A) or *gtrOC3* (D) mutations depends on capsule. Data presented as in Fig. 4 (n=3). (B) Liquid challenge assays. The indicated bacteria were cultured with or without Loki (MOI 1). Data are presented as in Fig 1A (n=3). (C) Plaque formation assays with Loki spotted on *A. baumannii* strains, and their corresponding *bfmS*-null derivative, harboring the indicated K locus.

**S7 Fig. Liquid phage challenge assays using *A. baumannii* strains BAA-1790 and 17978 and phages Loki or Loki*.** (A) The indicated BAA-1790 mutant, or WT control, was challenged with Loki. (B) 17978 WT was challenged with Loki, virulent Loki* derivative, or no phage control. Data points show geometric mean ± s.d. (n=3).

**S8 Fig. *A. baumannii* RnaA lacks catalytic residues defining the RNase T2 family.** Diagram shows analysis of conserved domains on RnaA. RNase_T2 domain, CAS motifs, and active site positions were identified using the NCBI Conserved Domain Database [87]. Signal peptide was identified using SignalP [88]. Vertical green lines represent conserved cysteine residues within the RNase_T2 domain. Alignment shows RNase T2 family homologs across diverse bacterial species. Within the conserved active site motifs (CAS-I and CAS-II), the active sites are indicated by #, and the most commonly found residues at these sites are highlighted.

**S9 Fig. Analysis of LOS production in altered phage susceptibility mutants bearing mutations in capsule-control loci or *lon*.** LOS production in cell lysates of mutants in *bfmRS* (A), *rnaA* and *dsbA* (B), and *lon* (C) was analyzed in comparison with WT control via Pro-Q 300 emerald staining. Representative gels are shown in top panels. Arrowhead indicates location of novel intermediate band associated with Δ*bfmS* (A). Quantification is shown in bottom panels. Bars show mean ± s.d. (n=3). Within each set of strains, no significant difference (P > 0.05) observed in total LOS levels by one-way ANOVA (A, B) or unpaired t-test (C).

**S10 Fig. One-step growth curves showing enhanced Loki multiplication in phage hypersensitive mutant hosts.** Data points show geometric mean PFU/ml ± s.d. (n=4). Infected cell number (used in burst size calculations) was approximately 1-1.5 x 10^4^ per ml.

**S1 Table. Strains, plasmids, and primers used in this study.**

**S2 Table. Spontaneous *gtr6* mutations conferring phage Loki resistance.**

**S1 Dataset. Loki challenge Tn-seq data.**

