## Supplementary material for "Genome-wide phage susceptibility analysis in *Acinetobacter baumannii* reveals capsule modulation strategies that determine phage infectivity": S1 Table

### S1 Table. Strains, plasmids, and primers used in this study

#### Strains

| Designation | Genotype or description | Strain ID | Reference |
|---|---|---|---|
| ***A. baumannii*** | | | |
| 17978 | cerebrospinal fluid isolate, AbaAL44$^+$ (“UN”) type, ATCC | EGA83 | [1, 2] |
| 17978 *bfmS** | ATCC 17978 *bfmS*(G467Dfs*19) | EGA127 | [3] |
| 17978 Δ*bfmS* | ATCC 17978 Δ*bfmS*::*aacC1* | EGA195 | [3] |
| 17978 Δ*bfmRS* | ATCC 17978 Δ*bfmRS*::*aacC1* | EGA495 | [7] |
| 17978 Δ*itrA* | ATCC 17978 Δ*itrA* | EGA295 | [3] |
| 17978 Δ*wzc* | ATCC 17978 Δ*wzc::aacC1*, isolate 27 | EGA106-27 | [3] |
| 17978 Δ*wzc wzc*$^+$ | ATCC 17978 Δ*wzc::aacC1*, *wzc*$^+$ by marker rescue of EGA106-27 | EGA260 | [3] |
| 17978 Δ*dsbA* | ATCC 17978 Δ*dsbA* | NRA134 | This work |
| 17978 Δ*lon* | ATCC 17978 Δ*lon* | JBA154 | This work |
| 17978 Δ*gtrOC3* | ATCC 17978 Δ*gtrOC3* | JBA160 | This work |
| 17978 Δ*gtrOC4* | ATCC 17978 Δ*gtrOC4* | JBA161 | This work |
| 17978 Δ*gtr6* | ATCC 17978 Δ*gtr6* | JBA202 | This work |
| 17978 Δ*rnaA* | ATCC 17978 Δ*rnaA* | JBA211 | This work |
| 17978 Δ*slt* | ATCC 17978 Δ*slt* | EGA520 | This work |
| 17978 Δ*rnaA*Δ*bfmRS* | ATCC 17978 Δ*rnaA* Δ*bfmRS* | JBA234 | This work |
| 17978 Δ*dsbA*Δ*bfmRS* | ATCC 17978 Δ*dsbA* Δ*bfmRS* | JBA235 | This work |
| 17978 *bfmS**Δ*itrA* | ATCC 17978 *bfmS*(G467Dfs*19) Δ*itrA* | JBA210 | This work |
| 17978 Δ*bfmS**Δ*gtr6* | ATCC 17978 Δ*bfmS*::*aacC1* Δ*gtr6* | JBA208 | This work |
| 17978 Δ*gtrOC3*Δ*itrA* | ATCC 17978 Δ*gtrOC3* Δ*itrA* | JBA212 | This work |
| 17978 Δ*gtrOC3*Δ*gtr6* | ATCC 17978 Δ*gtrOC3* Δ*gtr6* | JBA215 | This work |
| 17978 Δ*dsbA/-* | NRA134 with pYDE152 | JBA172 | This work |
| 17978 Δ*dsbA/dsbA* | NRA134 with pJE101 (P(IPTG)-*dsbA*) | JBA168 | This work |
| 17978 Δ*lon/-* | JBA154 with pYDE152 | JBA174 | This work |
| 17978 Δ*lon/lon* | JBA154 with pJE103 (P(IPTG)-*lon*) | JBA170 | This work |
| 17978 Δ*gtrOC3/-* | JBA160 with pYDE152 | JBA171 | This work |
| 17978 Δ*gtrOC3/ gtrOC3* | JBA160 with pJE100 (P(IPTG)-*gtrOC3*) | JBA167 | This work |
| 17978 Δ*gtrOC4/-* | JBA161 with pYDE152 | JBA173 | This work |
| 17978 Δ*gtrOC4/ gtrOC4* | JBA161 with pJE102 (P(IPTG)-*gtrOC4*) | JBA169 | This work |
| 17978 Δ*rnaA/-* | JBA211 with pYDE152 | JBA218 | This work |
| 17978 Δ*rnaA/-* | JBA211 with pJE127 (P(IPTG)-*rnaA*) | JBA217 | This work |
| 17978 Δ*itrA/-* | EGA295 with pYDE152 | JBA270 | This work |
| 17978 Δ*itrA/itrA* | EGA295 with pJE172 (P(IPTG)-*itrA*) | JBA271 | This work |
| 17978 + P(IPTG)-*rnaA* | ATCC 17978 WT with pJE127 (P(IPTG)-*rnaA*) | JBA266 | This work |
| 17978 *bfmS** + P(IPTG)-*rnaA* | EGA127 with pJE127 (P(IPTG)-*rnaA*) | JBA256 | This work |
| 17978 + P(IPTG)-*dsbA* | ATCC 17978 WT with pJE101 (P(IPTG)-*dsbA*) | JBA349 | This work |
| 17978 *gtr6*440$A_{9\rightarrow 8}$ | Loki non-susceptible derivative of WT ATCC 17978 (*gtr6*440$A_{9\rightarrow 8}$), 2 independent isolates | JBA149, JBA267 | This work |
| 17978 *wzx*377$T_{9\rightarrow 10}$ | Loki non-susceptible derivative of WT ATCC 17978 (*wzx*377$T_{9\rightarrow 10}$) | JBA150 | This work |
| 17978 *bfmS** *gtr6*440$A_{9\rightarrow 8}$ | Loki non-susceptible derivative of EGA127 (*gtr6*440$A_{9\rightarrow 8}$) | JBA151 | This work |
| 17978 *bfmS** *bfmR*(G100D) | Loki non-susceptible derivative of EGA127 [*bfmR*(G100D)] | JBA152 | This work |

### S1 Table (continued)

| | | | |
|---|---|---|---|
| 17978 *bfmS* bfmR*(T85I) | Loki non-susceptible derivative of EGA127 [*bfmR*(T85I)] | JBA153 | This work |
| 17978 *bfmS* gtr6*384$A_{8\rightarrow7}$ | Loki non-susceptible derivative of EGA127 (*gtr6*384$A_{8\rightarrow7}$) | JBA261 | This work |
| 17978 Δ*bfmS gtr6*440$A_{9\rightarrow8}$ | Loki non-susceptible derivative of EGA195 (*gtr6*440$A_{9\rightarrow8}$) | JBA268 | This work |
| 17978 *gtr6*440$A_{9\rightarrow8}$/- | JBA149 with pYDE152 | JBA204 | This work |
| 17978 *gtr6*440$A_{9\rightarrow8}$/*gtr6* | JBA149 with pJE121 (P(IPTG)-*gtr6*) | JBA205 | This work |
| 17978 *10840*p-GFP | ATCC 17978 WT with pEGE246 (*10840*p-GFP) | EGA616 | [7] |
| 17978 Δ*bfmRS 10840*p-GFP | EGA495 with pEGE246 (*18040*p-GFP) | NRA104 | This work |
| 17978 Δ*dsbA 10840*p-GFP | NRA134 with pEGE246 (*18040*p-GFP) | NRA178 | This work |
| 17978 Δ*dsbA*Δ*bfmRS 10840*p-GFP | JBA235 with pEGE246 (*18040*p-GFP) | NRA236 | This work |
| 17978 Δ*rnaA 10840*p-GFP | JBA211 with pEGE246 (*18040*p-GFP) | NRA238 | This work |
| 17978 Δ*rnaA*Δ*bfmRS 10840*p-GFP | JBA234 with pEGE246 (*18040*p-GFP) | NRA240 | This work |
| 17978 *adc*p-GFP | ATCC 17978 WT with pEGE313 (*adc*p-GFP) | EGA786 | [7] |
| 17978 Δ*dsbA adc*p-GFP | NRA134 with pEGE313 (*adc*p-GFP) | NRA237 | This work |
| 17978 Δ*rnaA adc*p-GFP | JBA211 with pEGE313 (*adc*p-GFP) | NRA239 | This work |
| 17961 | blood isolate, ATCC | EGA56 | [1] |
| 19606 | urine isolate, ATCC | EGA2M | [1] |
| 19606 Δ*bfmS* | ATCC 19606 Δ*bfmS*::*aacC1* | EGA216 | [3] |
| AB5075 | bone isolate/osteomyelitis, AB5075-UW | EGA714 | [4] |
| AB5075 *bfmS*::Tn | AB5075-UW *bfmS*115::T26 | EGA725 | [4] |
| BAA-1790 | sputum isolate (2008, Washington, DC), ATCC | JBA394 | [6] |
| BAA-1790 Δ*rnaA* | BAA-1790 Δ*rnaA*, via allele exchange with pJE193 | JBA395 | This work |
| BAA-1790 Δ*dsbA* | BAA-1790 Δ*dsbA*, via allele exchange with pJE195 | JBA396 | This work |
| EGA10 | wound isolate, Tufts University Medical Center, ST2 | EGA10 | This work |
| EGA65 | blood isolate, Tufts University Medical Center, ST2 | EGA65 | This work |
| EGA10/- | EGA10 with pYDE152 | JBA246 | This work |
| EGA10/*gtr6* | EGA10 with pJE121 (P(IPTG)-*gtr6*) | JBA247 | This work |
| EGA65/- | EGA65 with pYDE152 | JBA248 | This work |
| EGA65/*gtr6* | EGA65 with pJE121 (P(IPTG)-*gtr6*) | JBA249 | This work |
| ***E. coli*** | | | |
| DH5α | *supE44* Δ*lacU169* (ϕ80lacZΔM15) *hsdR17 recA1 endA1 gyrA96 thi*-1 *relA1* | EGE1 | [8] |
| DH5λpir | DH5α (λpir) *tet*::Mu *recA* | EGE4 | [9] |
| XL1-blue | *recA1 endA1 gyrA96 thi-1 hsdR17 supE44 relA1 lac* [F′ *proAB lacI*$^q$ZΔM15 Tn10 Tc$^r$)] | AFE81 | Stratagene |
| ***Bacteriophage*** | | | |
| Loki | vB_AbaS_Loki, lytic *Acinetobacter* siphovirus isolated from activated sewage sludge | Loki | [10] |
| Loki* | Derivative of Loki showing enhanced virulence vs 17978 and BAA-1790 | JBΦ-15C | This work |

### S1 Table (continued)

#### Plasmids

| Plasmid | Description | Reference |
|---|---|---|
| pUC18 | *oriColE1* MCS Cb$^r$ | [11] |
| pSR47S | Conditionally replicating allele exchange plasmid (*oriTRP4 oriR6K sacB* Km$^r$) | [12] |
| pJB4648 | Conditionally replicating allele exchange plasmid (*oriTRP4 oriR6K sacB* Gm$^r$) | [12] |
| pEGE305 | P(IPTG) shuttle vector (*ori-pBR322 ori*-pWH1277 *bla*::*lacI*$^q$-T5*lac*P Tc$^r$) | [7] |
| pYDE152 | P(IPTG) shuttle vector (*ori-pBR322 ori-pWH1277 bla*::*lacI*$^q$-T5*lac*P-MCS Tc$^r$) | [13] |
| pDL1100 | *Himar1 mariner* (Km$^r$) delivery plasmid, C9 transposase (*ori-pSC101* Cb$^r$) | [14] |
| pJE89 | pUC18 containing homology arm upstream of *gtrOC3* | This work |
| pJE90 | pUC18 containing homology arm downstream of *gtrOC3* | This work |
| pJE93 | pJB4648::Δ*gtrOC3* allele exchange construct | This work |
| pJE91 | pUC18 containing homology arm upstream of *gtrOC4* | This work |
| pJE92 | pUC18 containing homology arm downstream of *gtrOC4* | This work |
| pJE94 | pJB4648::Δ*gtrOC4* allele exchange construct | This work |
| pJE114 | pUC18 containing homology arm upstream of *gtr6* | This work |
| pJE115 | pUC18 containing homology arm downstream of *gtr6* | This work |
| pJE118 | pJB4648::Δ*gtr6* allele exchange construct | This work |
| pJE122 | pUC18 containing homology arm upstream of *rnaA* | This work |
| pJE123 | pUC18 containing homology arm downstream of *rnaA* | This work |
| pJE124 | pJB4648::Δ*rnaA* allele exchange construct | This work |
| pEGE242 | pSR47S::Δ*slt* allele exchange construct | This work |
| pEGE133 | pSR47S::Δ*bfmRS*::*aacC1* allele exchange construct | [3] |
| pEGE125 | pSR47S::Δ*bfmS*::*aacC1* allele exchange construct | [3] |
| pEGE181 | pSR47S::Δ*itrA* allele exchange construct | [3] |
| pJE95 | pUC18::*gtrOC3* | This work |
| pJE99 | pUC18::*gtrOC4* | This work |
| pJE120 | pUC18::*gtr6* | This work |
| pJE171 | pUC18::*itrA* | This work |
| pJE126 | pUC18::*rnaA* | This work |
| pJE97 | pUC18::*lon* | This work |
| pJE98 | pUC18::*dsbA* | This work |
| pJE100 | pYDE152::*gtrOC3* (P(IPTG)-*gtrOC3*) | This work |
| pJE102 | pYDE152::*gtrOC4* (P(IPTG)-*gtrOC4*) | This work |
| pJE121 | pYDE152::*gtr6* (P(IPTG)-*gtr6*) | This work |
| pJE172 | pYDE152::*itrA* (P(IPTG)-*itrA*) | This work |
| pJE127 | pYDE152::*rnaA* (P(IPTG)-*rnaA*) | This work |
| pJE103 | pYDE152::*lon* (P(IPTG)-*lon*) | This work |
| pJE101 | pYDE152::*dsbA* (P(IPTG)-*dsbA*) | This work |
| pEGE246 | *ACX60_RS18040*p-*gfpmut3* reporter plasmid (*ori-pBR322 ori*-pWH1277, Tc$^r$) | [7] |
| pEGE313 | *adc*p-*gfpmut3* reporter plasmid (*ori-pBR322 ori*-pWH1277, Tc$^r$) | [7] |
| pJE193 | pSR47S::Δ*rnaA*(BAA-1790) allele exchange construct | This work |
| pJE195 | pSR47S::Δ*dsbA*(BAA-1790) allele exchange construct | This work |

### S1 Table (continued)

#### Oligonucleotide primers

| Primer name | Sequence (5' – 3'; restriction site underlined if present) | RE site(s) |
|---|---|---|
| ***Complementation and localization experiments*** | | |
| dsbA-F | CGAGCTCTAGAGGAAAAGCTGTAACAATGAAA | SacI |
| dsbA-R | AAAACTGCAGGCAATAAACTATTATTTTGCCTTAC | PstI |
| lon-F | CGAGCTCATTAGGAGTGCCCATGTCTG | SacI |
| lon-R | AAAACTGCAGGTGAATTAGTGACGCGCTGCTTTTG | PstI |
| gtrOC3-F | CGAGCTCGCTAAAGGACGTTATAAGTTATGAAT | SacI |
| gtrOC3-R | AAAACTGCAGGCTTGCCTACTTATTTTTATCTTTATTC | PstI |
| gtrOC4-F | CGAGCTCGATAAAAATAAGTAGGCAAGCTATGAAAATTG | SacI |
| gtrOC4-R | AAAACTGCAGTAATTAAGTTTTCAGCAGGCCTTTAT | PstI |
| gtr6-F | CGAGCTCTAAGGTTACTATATGAAAATTGGATTG | SacI |
| gtr6-R | AAAACTGCAGCACCTACGACATCATTATTAGGTACA | PstI |
| itrA-F | CGAGCTCGACTGTTACCAGCGAATTTATTA | SacI |
| itrA-R | TGTACTGCAGCCGGCTACTGGTAAAACTG | PstI |
| rnaA-F | CGAGCTCAATGATGGAACATTCTTTTAATTG | SacI |
| rnaA-R | AAAACTGCAGGGTTATTGCTCTTAATAACTGCCT | PstI |
| ***Gene deletion*** | | |
| gtrOC4-upF | ACGCGTCGACGTGCCCGAGTTTTGCTTATC | SalI |
| gtrOC4-upR | GGGGTACCCTGAACAATTTTCATAGCTTGCCTAC | KpnI |
| gtrOC4-downF | GGGGTACCGGCCTGCTGAAAACTTAATTATTTTTTGGTG | KpnI |
| gtrOC4-downR | AAATATGCGGCCGCGTATATGCGCTTTGGCTGGT | NotI |
| gtrOC3-upF | ACGCGTCGACTGACTTGGCCAAAAAGATCC | SalI |
| gtrOC3-upR | GGGGTACCACATGGAATTTTATTCATAACTTATAACG | KpnI |
| gtrOC3-downF | GGGGTACCTAGGCAAGCTATGAAAATTGTTCAGGT | KpnI |
| gtrOC3-downR | AAATATGCGGCCGCTCAGTTCAGCCTTGGCTTTT | NotI |
| gtr6-UP-F | ACGCGTCGACGGCATATGCAATCTCAGG | SalI |
| gtr6-UP-R | GGGGTACCTGTCAATCCAATTTTCATATAGTA | KpnI |
| gtr6-DOWN-F | GGGGTACCACTATTGGTATTTAATATGTTATCTTGT | KpnI |
| gtr6-DOWN-R | AAATATGCGGCCGCATGATGCTAGCCCAATCC | NotI |
| Dlon-up-F | GGATCCTTAGCGCTTCCTTCACG | BamHI |
| Dlon-up-R | GGTACCCATAATAAGTTCAGACATGGGC | KpnI |
| Dlon-dwn-F | GGTACCGCAGCGCGTCACTAATTC | KpnI |
| Dlon-dwn-R | GTCGACCGATGTAAGGCAGGACCC | SalI |
| rnaA-UP-F | ACGCGTCGACGCGATAGTAATTCTAAAAACACG | SalI |
| DrnaA-UP-F1790 | ACGCGTCGACGCGATAGTAATTCTAAGAATACGG | SalI |
| rnaA-UP-R | GGGGTACCATATTTCATTTAATTAAAACTTGGTTATG | KpnI |
| rnaA-DOWN-F | GGGGTACCGGCAGTTATTAAGAGCAATAACCA | KpnI |
| rnaA-DOWN-R | AAATATGCGGCCGCAGAAGCAGCAGAAGACAAAC | NotI |
| Dslt-upF | CACCTGGATCCGTCTCACCACGATATCCATTCAC | BamHI |
| Dslt-upR | CATAGGTACCTTTCATCACACGACTCTTGTTTG | KpnI |
| Dslt-dwnF | AGTCGGTACCACGCGATCATCACCTTAGTTTTAG | KpnI |
| Dslt-dwnR | ATCTTGCGGCCGCATTTCACCGCTTGCGATAC | NotI |

### S1 Table (continued)

| | |
|---|---|
| ***qRT-PCR*** | |
| wza-F | CTGGCTATTCACTGCACAAAC |
| wza-R | CCTGACTCACCCATCACATAAA |
| gna-F | GCGGGTAGTAAGTGGAACTTT |
| gna-R | AACCTACCTCTTCTGCTTTATGG |
| rnaA-qRT-F | GTGCGCTAGACCCTTCTAAAC |
| rnaA-qRT-R | ACAGTCAGTTCTGGTGGTTTC |
