## Supplementary material for "Genome-wide phage susceptibility analysis in *Acinetobacter baumannii* reveals capsule modulation strategies that determine phage infectivity": S2 Table

**S2 Table. Spontaneous *gtr6* mutations conferring phage Loki resistance**

| *gtr6* mutation | residue changes | strain(s), this study | KL | additional strains (Genbank)[b] | | | additional strains (PATRIC)[b] | | |
|---|---|---|---|---|---|---|---|---|---|
| | | | | strain name | accession | KL | strain name | accession | KL |
| 440A$_{9\rightarrow8}$ | V142Gfs*8 | JBA149[a] | 3 | VB2107 | CP051474.1 | 3 | 2015ZJAB25 | WQMN01000013 | 3 |
| | | JBA151[a] | 3 | ABUH763 | CP035051.1 | 22 | 2087 | VMKD01000009 | 3 |
| | | JBA267[a] | 3 | AC29 | CP007535.2 | 3 | 208510 | VMJJ01000010 | 3 |
| | | JBA268[a] | 3 | AC30 | CP007577.1 | 3 | 24860_4 | JFDE01000009 | 22 |
| | | | | 2018BJAB1 | CP059351.1 | 3 | 4300STDY7045687 | UFIZ01000012 | 3 |
| | | | | 2018BJAB2 | CP059350.1 | 3 | 4300STDY7045767 | UFKE01000006 | 3 |
| | | | | 2022CK-00784 | CP117764.1 | 3 | Ab83 strain D86 | UEJK01000064 | 3 |
| | | | | | | | C248 | JAAZUG010000023 | 3 |
| | | | | | | | PWa17_1044 | JAJMQN010000011 | 3 |
| | | | | | | | PWb12_715 | JAJMPV010000012 | 3 |
| | | | | | | | PWb48_2085 | JAJMPN010000016 | 3 |
| | | | | | | | PWb63_2087 | JAJMPG010000011 | 3 |
| | | | | | | | SP2107 | JAAGSY010000002 | 3 |
| 384A$_{8\rightarrow7}$ | M150Cfs*22 | JBA261[a] | 3 | none | | | none | | |
| 425::IS*Aba13* | I131Ffs*6 | EGA10 | 22 | none | | | none | | |
| | | EGA65 | 22 | | | | | | |

[a] ATCC 17978 background

[b] listed genomes had >75x sequencing coverage except C248 (24x), and no assembly anomalies
